# For MSTd, Autoencoding is all you need

**DOI:** 10.64898/2026.03.23.713701

**Authors:** Oliver W. Layton, Scott T. Steinmetz

## Abstract

While goal-driven artificial neural networks (ANNs) have successfully modeled important aspects of the primate ventral stream, their efficacy for the dorsal stream remains unclear. Here, we investigated how computational objectives and architectural constraints influence the neural alignment to MSTd, a dorsal area that demonstrates selectivity to complex optic flow patterns and is linked to self-motion perception. We systematically evaluated the neural alignment between 54 ANNs and Non-negative Matrix Factorization (NNMF) against key neurophysiological optic flow tuning properties of MSTd. We optimized these models on either a supervised self-motion estimation task (accuracy-optimized) or an unsupervised input reconstruction task (autoencoding) using both raw optic flow and model MT-encoded signals. Interestingly, accuracy on the self-motion task does not predict neural alignment. Instead, model performance bifurcates based on both objective and input encoding: autoencoders utilizing MT-like input signals consistently achieve superior correspondence with MSTd tuning preferences. Explicitly enforcing sparsity or non-negativity does not improve alignment; rather, these constraints often degrade the match to biological data. Furthermore, we demonstrate that neural alignment remains largely unaffected even when the pressure to generate an efficient code with few units is eased, suggesting that dimensionality reduction may not be a primary driver of MSTd-like tuning. Taken together, our results indicate that the tuning properties of MSTd are better explained by an unsupervised reconstruction-based objective than by supervised task optimization, suggesting a fundamental difference in the computational principles that govern the dorsal and ventral streams.

**Significance Statement:** Goal-driven neural networks have revolutionized our understanding of the ventral visual stream, yet their effectiveness in modeling the dorsal stream remains less clear. We systematically evaluated 54 neural network models to identify the computational principles that drive neural-like optic flow tuning in dorsal stream area MSTd. Surprisingly, we find that accuracy-optimized models fail to replicate biological tuning. Instead, models that reconstruct motion inputs from a biologically plausible MT-like representation achieve the highest consistency with MSTd neurons. These findings suggest that the organizational principles of dorsal stream area MSTd may be better explained by an unsupervised, reconstruction-based objective rather than one focused on the accuracy of self-motion estimation, suggesting a fundamental difference in the computational objectives of the two visual streams.

## Introduction

The two-streams hypothesis revolutionized visual neuroscience by replacing unitary theories (Hubel and Wiesel, 1962; Marr, 1982) with a framework that emphasized functional organization (Ungerleider, 1982; Goodale and Milner, 1992): a ventral ‘what’ pathway that subserves object recognition (Dicarlo et al., 2012) and a dorsal ‘where’ pathway that mediates motion and action (Van Essen and Maunsell, 1983).

This functional focus has similarly transformed computational modeling. While traditional mechanistic models provided insights into neurophysiological plausibility (Fukushima, 1980; Riesenhuber and Poggio, 1999; Serre et al., 2007; Fazl et al., 2009), they struggle to capture the complexity of neural responses as effectively as modern artificial neural networks (ANNs) (Yamins et al., 2013; Khaligh-Razavi and Kriegeskorte, 2014; Lindsay, 2021). Rather than manually tuning parameters to mimic biological properties, ANNs offer a goal-driven framework where model representations emerge through the optimization of a specific objective (Rumelhart et al., 1986). Over the last decade, deep convolutional architectures trained on large-scale image classification—a task inherently aligned with the object recognition function associated with the ventral stream (Dicarlo and Cox, 2007)—have become the leading models of ventral stream neurons (Krizhevsky et al., 2012; Yamins et al., 2013; Simonyan and Zisserman, 2014; He et al., 2015). Notably, ANNs with higher classification accuracy on datasets like ImageNet tend to exhibit greater neural consistency (Schrimpf et al., 2020) (though see Linsley et al., 2024).

Inspired by the success of goal-driven modeling in the ventral stream, we investigated whether this approach extends to the dorsal stream, a question that has received less attention (Mineault et al., 2021; Vafaii et al., 2024; Layton and Steinmetz, 2024). We recently focused on ANNs optimized for self-motion estimation, a fundamental task ascribed to the dorsal stream (Saito et al., 1986; Duffy and Wurtz, 1991; Britten, 2008). Specifically, we explored whether these accuracy-optimized models could capture the motion tuning of neurons in the macaque medial superior temporal area (MSTd), a dorsal stream region that has been causally linked to self-motion perception (Gu et al., 2012). MSTd receives its primary afferent input from the middle temporal area (MT), where neurons are tuned for the local direction and speed of motion (Allman et al., 1985; Born and Bradley, 2005). MSTd exhibits tuning to full-field, complex patterns of retinal motion known as optic flow (Figure 1a–f), which arise during self-motion and provide rich information from which primates are capable of judging their trajectory (Warren et al., 1988; Britten and Van Wezel, 1998). While our ANNs successfully learned to estimate self-motion from optic flow, model units failed to capture key MSTd tuning properties (Layton and Steinmetz, 2024). By contrast, non-negative matrix factorization (NNMF), which performs unsupervised dimensionality reduction on motion signals, achieved superior neural consistency (Beyeler et al., 2016).

**Figure 1.**
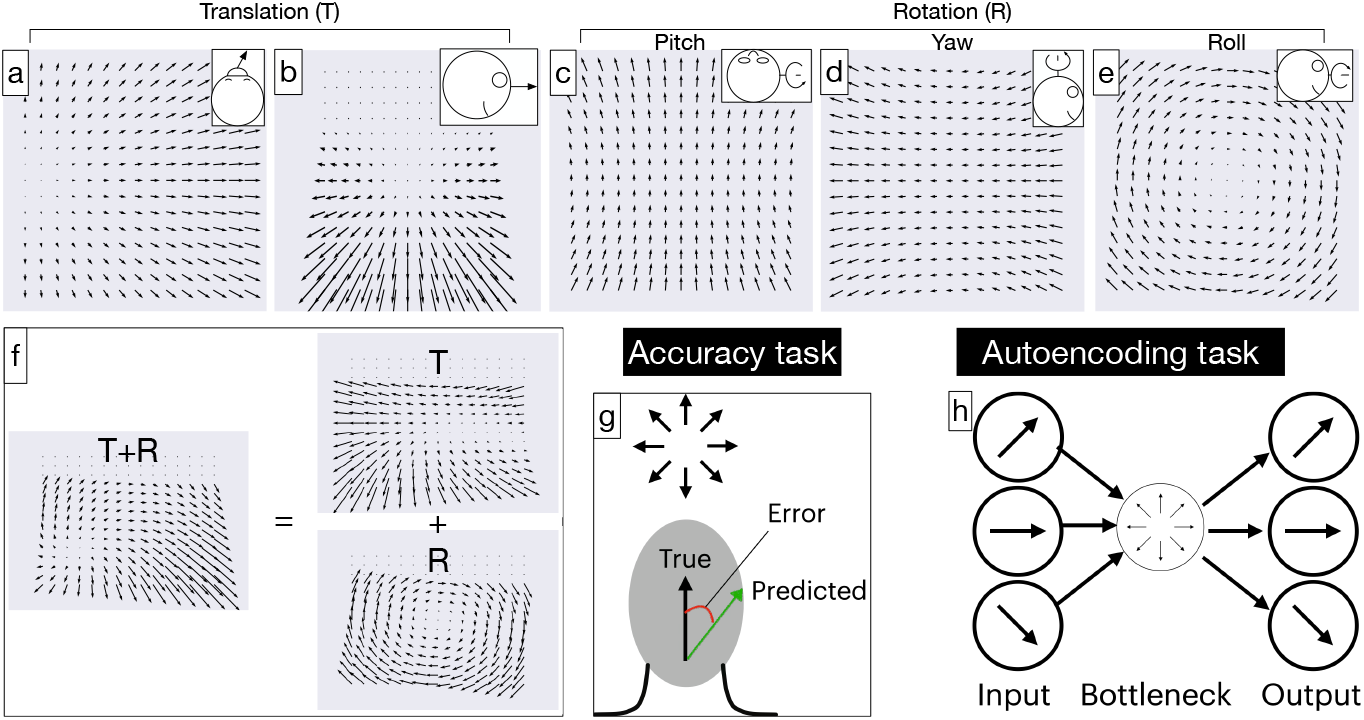
Optic flow field examples and computational modeling objectives. **a**, Optic flow produced by translational movement toward a frontoparallel wall (azimuth: 135°, elevation: 0°). **b**, Optic flow produced by straightforward translational movement above a ground plane with a 30° downward gaze offset (azimuth: 90°, elevation: 0°). **c**–**e**, Optic flow produced by pitch, yaw, and roll rotation about the *x*-, *y*-, and *z*-axes, respectively. **f**, Optic flow produced by a combination of translational (azimuth: 135°, elevation: 0°) and rotational movement (azimuth: 90°, elevation: 0°) above a ground plane with a 30° downward gaze offset. **g**, Schematic of accuracy-optimized ANNs trained to minimize the error between true and predicted self-motion directions. **h**, Schematic of autoencoder ANNs trained to minimize the error in reconstructing the optic flow signal from a latent representation.

In the present work, we investigated whether incorporating computational objectives and constraints inspired by NNMF yields more effective models of MSTd. NNMF is characterized by five core components: (1) an unsupervised reconstruction objective (informational bottleneck); (2) MT-like encoding of speed and direction rather than raw vector fields; (3) dense connectivity to facilitate local motion integration; (4) non-negative, linear activations; and (5) non-negative weights. These factors diverge substantially from effective ANN models of the ventral stream, which typically utilize supervised learning, raw image rather than neurally embedded inputs (e.g. MT activations), convolutional connectivity, non-linear integration, and unconstrained weight signs. To isolate the contribution of each factor, we trained 54 ANNs across various combinations of these constraints and quantitatively compared their optic flow tuning to MSTd data.

Our simulations reveal that linear autoencoders—ANNs implementing the NNMF reconstruction objective without the non-negativity constraints—yield the highest consistency with MSTd, performing comparably to the original NNMF model. We found that shallow rather than deep architectures better supported neural alignment, as did learning from an MT-encoded motion signal rather than from raw optic flow.

## Materials and Methods

### Datasets

We trained ANNs on a dataset composed of 6,030 optic flow fields of size 15*×*15, generated from random combinations of simulated 3D translational and rotational components of self-motion. We refer to this dataset as TR360, which has been used in modeling studies of MSTd (Beyeler et al., 2016; Layton and Steinmetz, 2024; Layton et al., 2024). Each optic flow vector 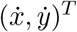 at position (*x, y*) was computed using the standard optic flow equations for a pinhole camera model (Longuet-Higgins and Prazdny, 1980; Raudies and Neumann, 2013):

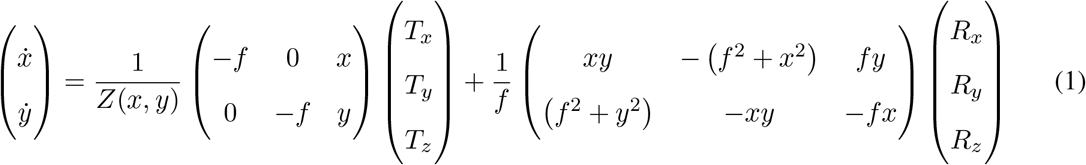

In Eq. 1, 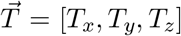 denotes the 3D translational velocity of the observer, 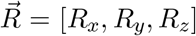 denotes the 3D rotational velocity of the observer, *f* is the focal length of the camera model (fixed at 1 cm), and *Z*(*x, y*) is the depth of the projected point in the environment at position (*x, y*). Following Beyeler et al. (2016), half of the dataset corresponds to movement toward a frontoparallel wall (e.g. Figure 1a), and the other half corresponds to movement over a ground plane with a 30° downward gaze offset (e.g. Figure 1b). To simulate the ground plane scenario, we set 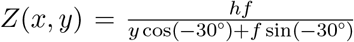, where *h* = −10 m corresponds to the observer’s eye height (Beyeler et al., 2016). We zero padded the optic flow fields spatially to 16*×*16 before presenting the data samples to the ANNs to permit more consistent resizing throughout network layers. Table 1 summarizes the parameters used to generate the TR360 dataset.

**Table 1.** Optic flow datasets used for model training and evaluation. TR denotes optic flow generated from combined translational (*T*) and rotational (*R*) self-motion. Only the *T R*360 dataset was used for ANN training; all other datasets served as test sets for characterizing optic flow tuning properties. Straight-ahead heading corresponds to an azimuth of 90° and an elevation of 0°.

| Dataset | Description | Size (N samples) | Independent variables |
| --- | --- | --- | --- |
| **TR360** | Simulated self-motion toward either a frontoparallel plane (no gaze offset) or a ground plane ($-30^\circ$ gaze offset). Translation and rotation directions drawn uniformly at random: TR elevation $[0, 360]^\circ$, TR azimuth $[90, -90]^\circ$. | Total: 12,060 (6,030 wall + 6,030 ground)<br>Train: 6,030<br>Validation: 3,015<br>Test: 3,015 | T speed: $[0.5, 1.0, 1.5]$ m/s<br>R speed: $[0, 5, 10]^\circ/\text{s}$<br>Plane depth: $[2, 4, 8, 16, 32]$ m |
| **BenHamedT** | Translational optic flow fields used to compare ANN decoding accuracy with that achieved by Ben Hamed et al. (2003). 3D dot cloud simulated environment (depth: 1–8 m). T azimuth and elevation randomly selected at $30^\circ$ eccentricity. | 1,500 samples | T speed: $[0.5, 2.0]$ m/s |
| **TestProtocolT** | Diagnostic set used to evaluate tuning to specific translation directions. | 497 samples (495 azimuth/elevation combinations + $\pm 90^\circ$ vertical) | T azimuth: $0, 11.25, 22.5, \dots, 360^\circ$<br>T elevation: $0, \pm 11.25, \pm 22.5, \dots, \pm 90^\circ$ |
| **TestProtocolR** | Diagnostic set used to evaluate tuning to specific rotation directions. | 497 samples (495 azimuth/elevation combinations + $\pm 90^\circ$ vertical) | R azimuth: $0, 11.25, 22.5, \dots, 360^\circ$<br>R elevation: $0, \pm 11.25, \pm 22.5, \dots, \pm 90^\circ$ |

### MT encoding of optic flow

Whereas some ANNs were trained directly on the raw TR360 optic flow vectors, others were trained on motion signals generated by a computational model of area MT (Beyeler et al., 2016). This model represents MT speed tuning using a log-normal distribution (Nover et al., 2005) and direction tuning using a von Mises function to capture the falloff in responses as stimulus direction deviates from the preferred direction (Allman et al., 1985). Specifically, the speed tuning of a model MT unit with a receptive field is centered at (*x, y*) that preferentially responds to speed *ν*_*pref*_ obeys

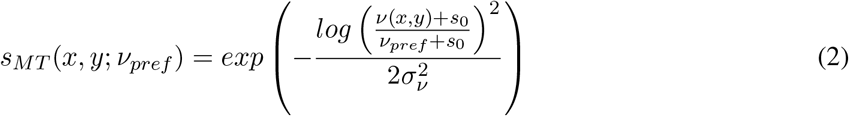

where *ν*(*x, y*) is the speed of the optic flow vector at position (*x, y*) and *s*_0_ is a small positive constant that ensures that logarithm is well-defined. We set *σ*_*ν*_ = 1.16 and *s*_0_ = 0.33 to match median values fitted from MT neuron recordings (Nover et al., 2005).

We modeled MT direction tuning using a von Mises distribution so that responses fall off symmetrically as the stimulus direction deviates from each unit’s preferred direction:

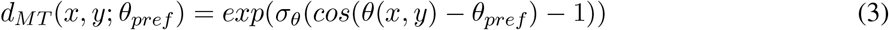

In Eq. 3, *θ*(*x, y*) denotes the direction of the optic flow vector at position (*x, y*), *θ*_*pref*_ is the preferred direction of the simulated neuron, and *σ*_*θ*_ controls the tuning width. We fixed *σ*_*θ*_ at 3° to produce a full-width at half-maximum tuning of approximately 90°, consistent with MT physiology (Britten and Van Wezel, 1998).

Each model MT unit integrated a single optic flow vector within the 15×15 optic flow field, yielding a receptive field of roughly 6° of visual angle under a 90° field of view. At each spatial position (*x, y*), we simulated 40 MT units with randomly assigned preferred speeds (2–32°/s) and directions, producing a 15×15×40 population. We multiplied the speed (Eq. 2) and direction (Eq. 3) tuning functions together to generate the response of each MT neuron:

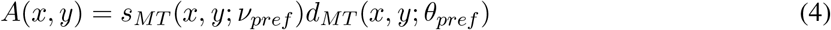

### Artificial neural networks

We trained ANNs on the TR360 optic flow dataset to optimize either the fidelity with which they reconstruct motion signals (autoencoders) or the accuracy with which they estimate the observer’s translational and rotational self-motion (accuracy-optimized ANNs). For the autoencoders, the training objective was to minimize the mean squared error (MSE) between the true and reconstructed optic flow vectors or MT unit activations, depending on whether the autoencoders operated on the raw optic flow or the MT encoded signal, respectively. The accuracy-optimized ANNs minimized the mean squared error (MSE) between each true and predicted self-motion component.

We parameterized rotational velocity using the 3D Cartesian vector 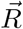 and translational velocity using spherical azimuth (*t*_*az*_) and elevation (*t*_*el*_) angles, consistent with conventions in the optic flow literature (Warren et al., 1988; Perrone, 2018). Separate output units estimated each of these five components. To account for the circularity of translational azimuth (0°–360°), the ANNs minimized a cosine-based loss rather than MSE:

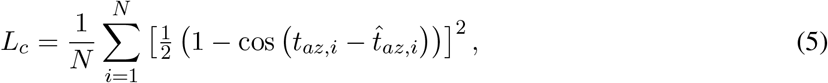

where *N* is the number of optic flow samples in the training mini-batch, *t*_*az,i*_ is the true azimuth of optic flow sample *i* and 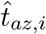 is the network’s estimate. Unlike MSE, the loss function *L*_*c*_ assigns the same penalty to a fixed angular error regardless of whether it crosses the circular boundary. For example, estimating 0° azimuth as 20° incurs the same loss as estimating it as 340°.

We constructed variants of the accuracy-optimized and autoencoder ANNs that differ in their architectural choices and mechanisms along the following dimensions:

- *Dense vs. convolutional connectivity:* In some networks, neurons in successive layers are fully connected (Figure 2a,c). In others, neurons in the early layers integrate inputs within local convolutional windows (Figure 2b,d).
- *Linear vs. ReLU activation:* Hidden layers either use a linear (identity) activation or a ReLU non-linearity in both convolutional (Conv2D) and fully connected (Dense) layers. The ReLU enforces non-negative neural responses, similar to the NNMF algorithm.
- *Non-negative weight constraint:* In a subset of models, we constrained all learned weights to be non-negative during training, consistent with the NNMF algorithm.
- *Network depth*: For each architecture, we trained shallow, medium-depth, and deep versions by varying the number of hidden layers.

**Figure 2.**
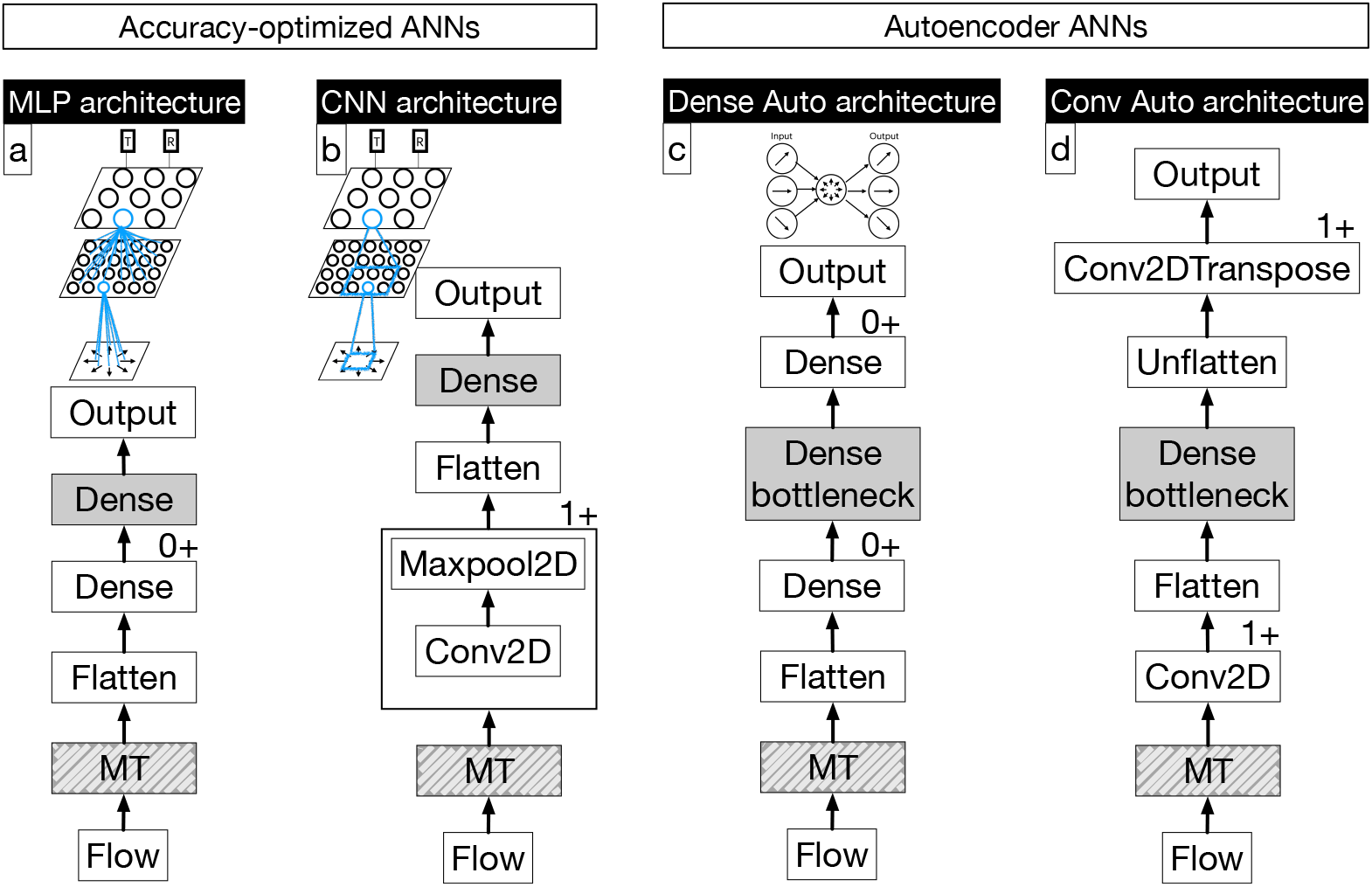
Schematics of the accuracy-optimized and autoencoder ANN architectures. **a, b**, Layout of the multi-layer perceptron (MLP) and convolutional neural network (CNN) accuracy-optimized architectures. **c, d**, Layout of the dense and convolutional autoencoder architectures, respectively. Models with “mt” in the name include the additional MT motion encoding layer (dashed box). Labels 0+ and 1+ indicate optional or repeated modules, respectively, whose counts vary across model variants. Hidden layers used for neural activation readout and subsequent alignment analysis appear in solid gray.

We followed several guiding principles when selecting hyperparameters for the ANNs to facilitate comparisons within and across architectures (Table 2). First, we aimed to equate the number of neurons in each readout layer, from which model activations were recorded and analyzed to assess optic flow tuning. This readout layer corresponds to the last hidden layer in accuracy-optimized ANNs and the middle “bottleneck” dense layer in the autoencoders (Figure 2). For most ANNs, we selected 1,024 neurons in this layer to reduce dimensionality from the 9,000 MT units (15*×*15*×*40) providing afferent motion inputs to the ANN.

**Table 2.**
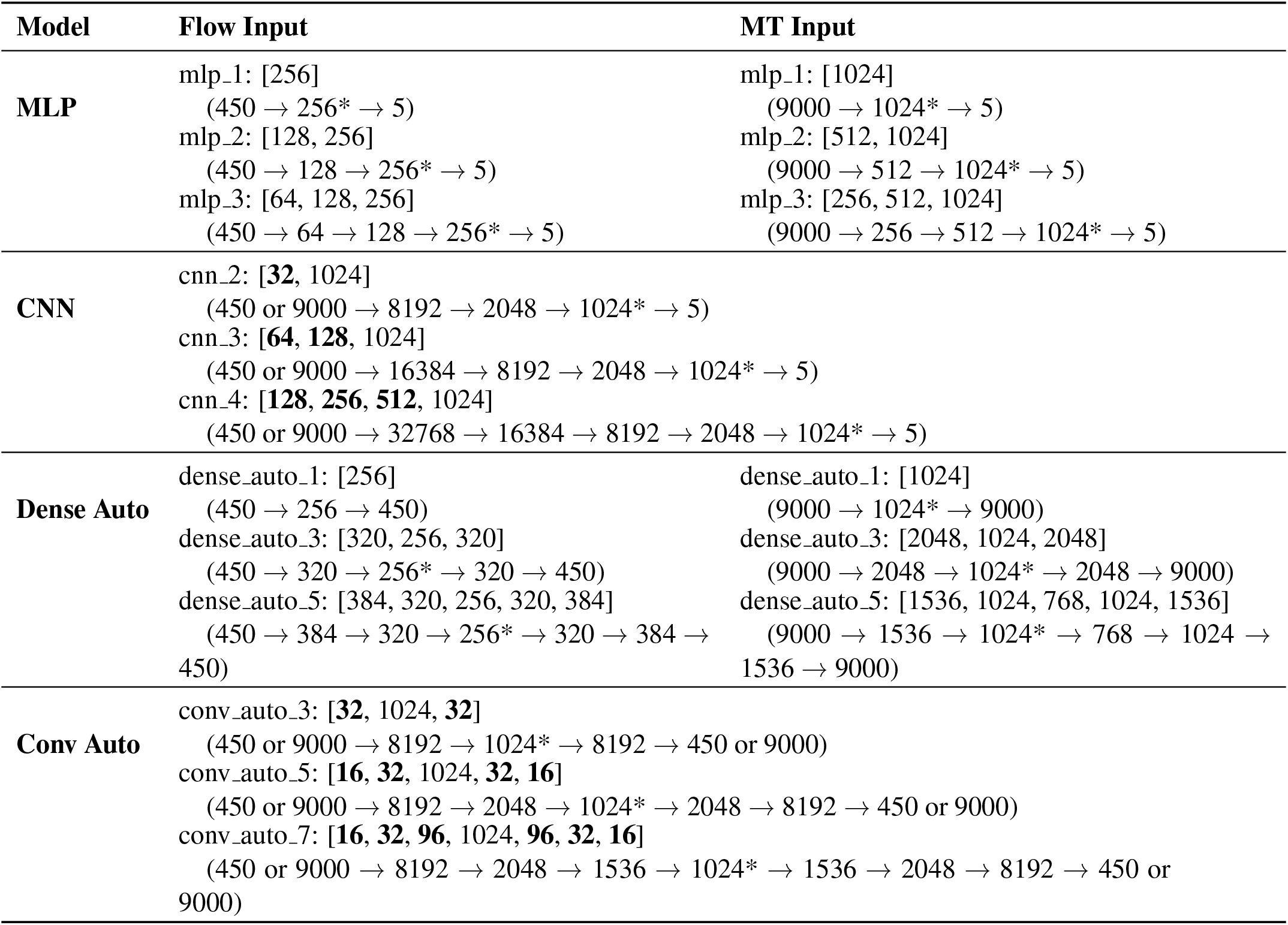
Configurations and layer dimensions of the 54 artificial neural networks. MLP and CNN architectures represent accuracy-optimized networks, while dense and convolutional variants represent autoencoders. The Flow Input column details configurations for raw optic flow (15 × 15 × 2), and the MT Input column lists those for MT-encoded inputs (15 × 15 × 40). We trained three variants of each model: shallow, medium, and deep. Square brackets contain the unit counts for each hidden layer (e.g., [128, 256] indicates 128 units in the first hidden layer and 256 in the second), with bold numbers denoting convolutional layers. Parentheses contain the total number of activations in each layer, and asterisks (∗) mark the readout layers used for neural alignment analysis.

This choice aligns with typical autoencoder design and the hypothesis that dimensionality reduction plays a key role in the visual system (Ballard, 1987; Ng, 2011; Beyeler et al., 2016, 2019). At the same time, 1,024 units provide a sufficiently large sample to reliably quantify population-level optic flow tuning. In our testing, altering the number of units in the readout layer did not qualitatively alter population tuning patterns, though decreasing it resulted in greater variability across training sessions.

In a limited number of networks, we were required to deviate from the target number of hidden units. Within non-convolutional ANNs trained on vector field representations of optic flow (MLP and dense autoencoder families), 1,024 neurons in the readout layer would not implement the desired dimensionality reduction because the input layer contains only 450 neurons (15*×*15*×*2). For these ANNs, we instead used 256 in the readout layer to reduce dimensionality while maintaining a reasonably large neural sample. In addition, the deepest dense autoencoder with 1,024 neurons in the bottleneck layer (dense auto 5 linear mt) failed to train due to memory limitations on our NVIDIA GeForce RTX 4090 24 GB GPU. We resolved this issue by reducing the bottleneck neurons slightly to 768.

Second, we configured the autoencoders to progressively reduce the dimensionality of motion signals while following the common ANN design principle of increasing the number of neurons in deeper layers. The spatial resolution of motion signals decreased concurrently in the convolutional ANNs.

Third, within the accuracy-optimized ANNs, we implemented the common design pattern in deep learning of progressively increasing the number of neurons from the first hidden layer onward. In the MLPs, but not the CNNs, this produced an initial decrease in dimensionality, followed by a progressive increase in the deeper layers. Nevertheless, the dimensionality of the readout layer is always lower than that of the input optic flow signal. We do not believe this design pattern impaired the alignment of these ANNs with neurophysiological properties. On the contrary, some of these MLPs garnered favorable correspondence across the optic flow tuning metrics that we evaluated.

In addition to the ANNs listed in Table 2, we trained two additional variants of dense auto 1 with MT-encoded inputs that impose a sparsity constraint, implemented in one of two ways. In one version (dense_ - auto_sparse_l1_1), we encouraged sparse coding through L1 regularization applied to the activations in each hidden (Tibshirani, 1996). We used a regularization strength of 10^−6^, as experimentation showed that this value produced the closest neural alignment with MSTd. In the second version (dense_auto_sparse_kl_1), we configured the network as a sparse autoencoder (Ng, 2011), applying a KL divergence penalty and using sigmoid instead of tanh activation functions throughout the network. We configured the KL divergence to promote a activation sparsity of 0.1. We scaled the penalty with a regularization strength of 0.01.

We trained all ANNs using the AdamW optimizer with early stopping (patience: 3 epochs) in TensorFlow 2.19. We used a batch size of 128 samples, Glorot uniform initialization for the weights, and zero initialization for the biases. We trained all ANNs with a learning rate of 0.001, except for conv auto 5 and conv auto 7 for which we used 0.00001. We configured the CNNs to possess 2×2 convolutional filters and 2×2 max pooling windows (stride: 2). For the convolutional autoencoders we used 3×3 convolutional filters with a stride of 2.

### Non-negative matrix factorization model

For comparison with the ANNs, we fitted the non-negative matrix factorization model (NNMF) of MSTd proposed by Beyeler et al. (2016) to the TR360 dataset. This model decomposes the MT activation matrix, *A*, into the product of two non-negative matrices, *H* and *W* :

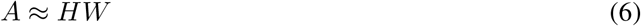

The MT activation matrix has shape (*N, M*), where *N* is the number of optic flow samples and *M* is the number of MT units. The matrix *H* has shape (*N, K*) and represents the activations of *K* = 1024 model MSTd units for each sample. The matrix *W* has shape (*K, M*) and encodes the weights between the *M* = 9000 MT units (15*×*15*×*40) and the *K* MSTd units. NNMF minimizes the mean squared error (MSE) reconstruction loss, *L*_*nnmf*_, between the reconstructed activations, *A*_*rec*_ = *HW*, and the original matrix, *A*:

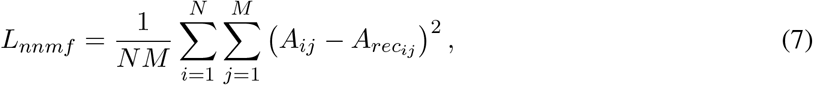

subject to the non-negativity constraints, *H*_*ik*_ ≥ 0 and *W*_*kj*_ ≥ 0. We implemented NNMF in TensorFlow and minimized *L*_*nnmf*_ using gradient descent with the Adam optimizer (learning rate: 0.01). Non-negativity was enforced at each training epoch via the ReLU function.

NNMF involves several hyperparameters. First, we initialized the entries of the matrices *H* and *W* from a uniform distribution between 0 and 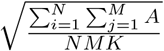, following Scikit-learn (Pedregosa et al., 2011). Second, the number of basis vectors, *K*, was determined by repeatedly fitting NNMF 16 times with 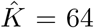 basis vectors per fit, then concatenated the resulting matrices to yield *K* = 1024. This procedure captures variability arising from random initialization in the fit (Beyeler et al., 2016). Third, we stopped training when the absolute change in loss between successive epochs was below 1 *×* 10^−4^, with a minimum of 2 epochs, consistent with the defaults of MATLAB’s `nnmf` function (Beyeler et al., 2016). This prevents overfitting during the iterative optimization of Eq. 7. Fourth, we used gradient descent for fitting, which aligns with our neural network training paradigm and leverages GPU acceleration. We obtained similar results when using the alternating least-squares algorithm employed by Beyeler et al. (2016).

Once fitted, MSTd activations for *N*_*test*_ novel stimuli were computed as:

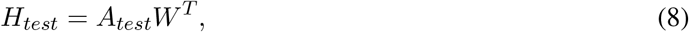

where *A*_*test*_ is the MT activation matrix for the test stimuli (shape: (*N*_*test*_, *M*)) and *W* ^*T*^ is the transpose of the fitted MT–MSTd basis vectors (shape: (*M, K*)).

### Optic flow tuning metrics

We characterized the optic flow tuning of individual units in the readout layer of each ANN using key metrics established in neurophysiological studies of MSTd:

- Preferred azimuth and elevation angles for pure translational and rotational self-motion (Takahashi et al., 2007; Gu et al., 2010)
- Difference between translational and rotational self-motion preferences (Takahashi et al., 2007)
- Heading sensitivity (Gu et al., 2010)
- Heading tuning index (HTI) (Gu et al., 2006)

To determine each model unit’s preferred translation direction, we presented each network with optic flow fields corresponding to pure translational self-motion. We systematically varied the horizontal and vertical directions in steps of 11.25° (TestProtocolT dataset in Table 1). We recorded the activations to each optic flow field and decoded the preferred 3D translation direction using the population vector approach (Georgopoulos et al., 1986):

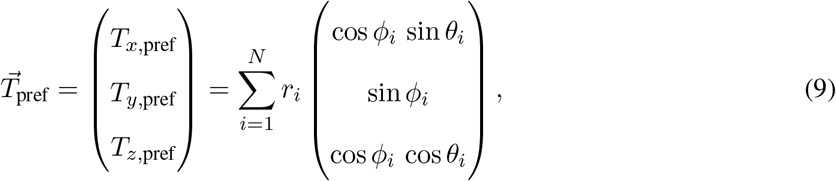

The vector 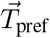 represents the estimated preferred 3D translation direction obtained by summing the unit responses weighted by their direction vectors. In Eq. 9, *r*_*i*_ denotes the response of the unit to optic flow field *i*, corresponding to self-motion along azimuth *θ*_*i*_ and elevation *ϕ*_*i*_. When computing the population vector estimate, we used a coordinate system in which *θ* spans [−180°, 180°] and *ϕ* spans [−90°, 90°]. We then converted 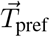 to the spherical coordinate system of Takahashi et al. (2007) (azimuth: [0°, 360°], elevation: [90°, −90°]) to allow comparison with MSTd neurons. We used the same procedure to determine each unit’s preferred direction of rotation using the diagnostic rotation dataset (TestProtocolR dataset in Table 1).

In addition to determining each unit’s preferred self-motion direction, we characterized its sensitivity to different horizontal translation directions (i.e. headings). This measure captures how strongly a unit’s activation changes in response to small deviations in heading. To facilitate comparison with the MSTd population data of (Gu et al., 2010), we quantified the heading that produced the largest change in activation (i.e. peak discriminability). To compute this, we presented each ANN with 24 optic flow patterns with headings spanning 0–360° (15° increments) and recorded the activations in each readout unit. We then fitted a cubic spline to the activation profile to interpolate smoothly between sampled headings. The heading corresponding to the maximum of the derivative of this spline was identified as the peak discriminability. To avoid artifacts at the circular boundary (0°/360°), the activation profile was circularly extended by repeating values across an additional 300° on either side before fitting the spline.

We also assessed the strength each readout unit’s heading tuning by computing the heading selectivity index (HTI). This metric ranges from 0 and 1, where 0 indicates a uniform response across all headings (i.e. poor selectivity) and 1 indicates high sensitivity for a narrow range of headings (i.e. strong selectivity).

Using the TestProtocolT diagnostic dataset, we computed the HTI for each model unit *j* according to Beyeler et al. (2016); Gu et al. (2006):

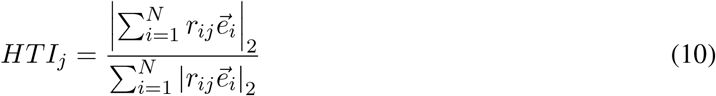

where *r*_*ij*_ denotes the activation of the *j*^th^ model unit in response to the *i*^th^ optic flow pattern, 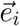 is the heading direction of the *i*^th^ stimulus expressed as a 3D Cartesian unit vector, and |·|_2_ denotes the *L*^2^ Euclidean norm.

### Neural alignment

We evaluated the extent to which units within the readout layer of each ANN develop optic flow tuning properties that align with those of MSTd neurons in neurophysiological experiments. In most analyses, we compared histograms summarizing population-level tuning patterns in ANN and MSTd neurons. We used the earth mover’s distance (EMD) to quantify the similarity between the normalized MSTd optic flow tuning metric histogram 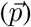 and that produced from the ANN population readout 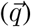:

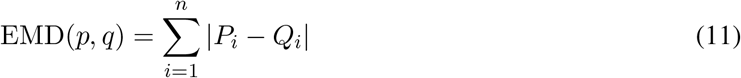

where *n* is the number of histogram bins, and *P*_*i*_ and *Q*_*i*_ are the respective cumulative distribution functions (CDFs) defined as:

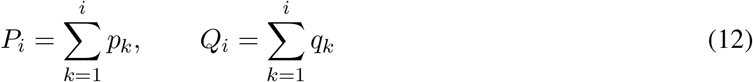

We selected EMD over KL divergence and other distance metrics because it accounts for transpositions of probability mass among nearby histograms bins. This is unlike KL divergence, which operates on a bin-by-bin basis and is insensitive to how far probability mass shifts; two distributions that differ only by small or moderate shifts may receive similar KL penalties. By contrast, EMD measures the cumulative discrepancy between CDFs and captures the cost of transporting mass between bins. This allows EMD to tolerate cases where ANN and MSTd histograms are largely similar, but exhibit small offsets in mass among nearby bins.

To assess to reliability of the EMDs for the optic flow tuning properties, we trained each of the 55 ANNs and NNMF 25 times with different random initial weights. We report the mean EMD and 95% confidence intervals computed across these 25 training repetitions.

Equation 11 computes the EMD for a single optic flow tuning property (e.g., preferred translation azimuth). To quantify overall neural alignment, we calculated a geometric mean of the EMDs across all tuning metrics for each ANN. Prior to this, we arithmetically averaged the EMDs across 25 training repetitions of each ANN, yielding 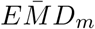 for model *m*. Because the raw EMD values differ in scale across tuning properties (due to differences in histogram bin size and other factors), we normalized 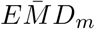 by the corresponding value from a baseline model (dense_auto_1_linear_mt). This normalization removes unit dependencies and allows the geometric mean to be computed consistently across tuning metrics.

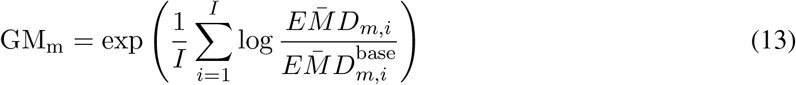

Eq. 13 quantifies the geometric mean for ANN model *m* across the *I* tuning metrics relative to the baseline model. Models with *GM <* 1 exhibit greater neural alignment than the baseline, whereas *GM >* 1 indicates weaker relative alignment. We compute the geometric mean in logarithmic space to improve numerical stability.

Because Gu et al. (2006) report the HTI for their MSTd population as a single point estimate rather than a distribution, we used the mean absolute error (MAE) to assess similarity to the ANNs. To reduce the influence of skewed distributions and units with extreme HTIs, we computed the MAE using the median HTI across all units in the readout layer of each ANN.

### Sparsity metrics

To quantify the sparsity of hidden layer representations, we employed population and lifetime metrics (Vinje and Gallant, 2000). Both measures stem from the following index:

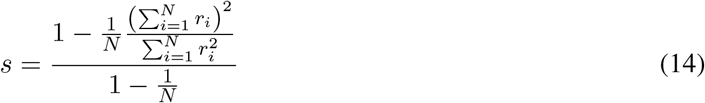

The index in Eq. 14 ranges from 0 to 1, where values approaching 1 represent a sparse code and values near 0 reflect a dense representation. We calculated these metrics as follows:

- *Population Sparsity:* In this context, *r*_*i*_ denotes the response of the *i*^*th*^ unit to a single optic flow stimulus, and *N* represents the total number of units in the layer. This metric quantifies the extent to which the network utilizes only a small subset of available units to encode any given motion pattern.
- *Lifetime Sparsity:* Here, *r*_*i*_ represents the response of a single unit to the *i*^*th*^ optic flow pattern in the dataset, and *N* is the total number of samples (3,015). This metric measures the selectivity of individual units over samples; a high value indicates that a unit responds only to a rare subset of optic flow fields.

We averaged the population sparsity across all stimuli and the lifetime sparsity across all units to produce single estimates for each model.

## Results

The success of goal-driven ANNs in the ventral stream inspired us to explore their efficacy in modeling MSTd optic flow tuning. Leading ventral models utilize supervised learning for image classification, so we optimized ANNs on an analogous dorsal task: self-motion estimation from optic flow. The objective of these “accuracy-optimized” models is to minimize the difference between true and estimated translational (T) and rotational (R) directions (Figure 1f,g). Given that NNMF successfully models optic flow tuning in MSTd (Beyeler et al., 2016), we also designed ANNs that implement its defining computational mechanisms. This includes NNMF’s unsupervised computational objective: reconstructing motion inputs through an informational bottleneck (Figure 1h). These “autoencoder” ANNs (Rumelhart et al., 1986; Ballard, 1987) are more biologically plausible than the accuracy-optimized objective because they do not require self-motion labels during learning (i.e. 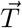 and 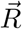).

To discover the factors driving neural consistency, we designed 54 models crossing the objectives and constraints of both NNMF and traditional ANNs. All models were trained on the TR360 dataset (Beyeler et al., 2016; Layton and Steinmetz, 2024), which consists of optic flow generated by combinations of translational and rotational optic flow (T+R; Figure 1f). To readily identify each model based on its defining characteristics, we created a model naming scheme (e.g. *dense_auto_1_linear_mt*) based on five architectural and functional factors:

1. *Objective and Connectivity (cnn, mlp, dense_auto, conv_auto): cnn* and *mlp* denote accuracy-optimized models using AlexNet-like convolutional (Krizhevsky et al., 2012) or multi-layer perceptron-like dense connectivity, respectively. *dense_auto* and *conv_auto* denote autoencoders with dense (NNMF-like) or convolutional connectivity.
2. *Depth (1–7):* The number of hidden layers that contain trainable weights. We tested three variants of each architecture (Figure 2) to contrast the shallow, single hidden layer structure of NNMF with the deep networks that model the ventral stream.
3. *Activation function (linear, relu):* Models use either linear activations (NNMF-like) or the nonlinear ReLU function common in ventral ANNs. Note that the CNN architectures are inherently nonlinear due to their max pooling stages.
4. *Input motion encoding (flow or mt)*: Models received either raw optic flow vectors (*flow*) or the population activations of 9,000 model MT neurons (*mt*) that encode those patterns.
5. *Weight constraint (nonneg)*: When present, the training protocol enforces the non-negative synaptic weights characteristic of NNMF.

In addition to the models described above, we developed two specialized variants of the single-layer dense autoencoder to evaluate the role of sparsity, a principle previously linked to optic flow tuning in MSTd (Beyeler et al., 2016, 2019). The first variant, *dense_auto_1_sparse_kl_mt*, enforces sparsity via a KL-divergence penalty with a target mean activation (*ρ*) of 0.1 for the hidden units (Ng, 2011; Goodfellow et al., 2016). This value was chosen empirically to consistently induce a sparse bottleneck representation. The second variant, *dense_auto_1_sparse_l1_mt*, utilizes L1 regularization on the bottleneck layer hidden activations to induce sparsity. These models allow us to determine whether explicitly enforced sparsity, independent of non-negativity constraints, improves neural consistency with MSTd.

### Task performance

To evaluate task performance, we compared the accuracy of the direction of translation (heading) decoded from ANN hidden layers with that obtained from a MSTd population. When linearly decoding MSTd responses to translational optic flow (30° heading eccentricity), Ben Hamed et al. (2003) found single-trial mean absolute errors (MAEs) of 1.57 ± 3.04° SD and 0.63 ± 2.39° SD for horizontal (x) and vertical (y) heading components, respectively. We performed an analogous analysis by fitting linear regressions (10-fold cross validation) to the hidden layer activations of each ANN using a similar dataset. For the accuracy-optimized ANNs, we used the final hidden layer and for the autoencoders we used the middle bottleneck layer.

As shown in Figure 3a, model performance bifurcates according to input signal encoding: ANNs learning from MT-encoded signals achieve the lowest decoding errors, many at least as low as those obtained from MSTd, whereas those processing raw optic flow yield substantially higher errors. This division persists largely regardless of computational objective, connectivity, or depth. These results suggest an intrinsic advantage of the MT motion representation for supporting accurate self-motion decoding, a representation that even the deeper models do not apparently learn from task constraints alone. It is noteworthy that the NNMF decoding accuracy is comparable to leading ANN models.

**Figure 3.**
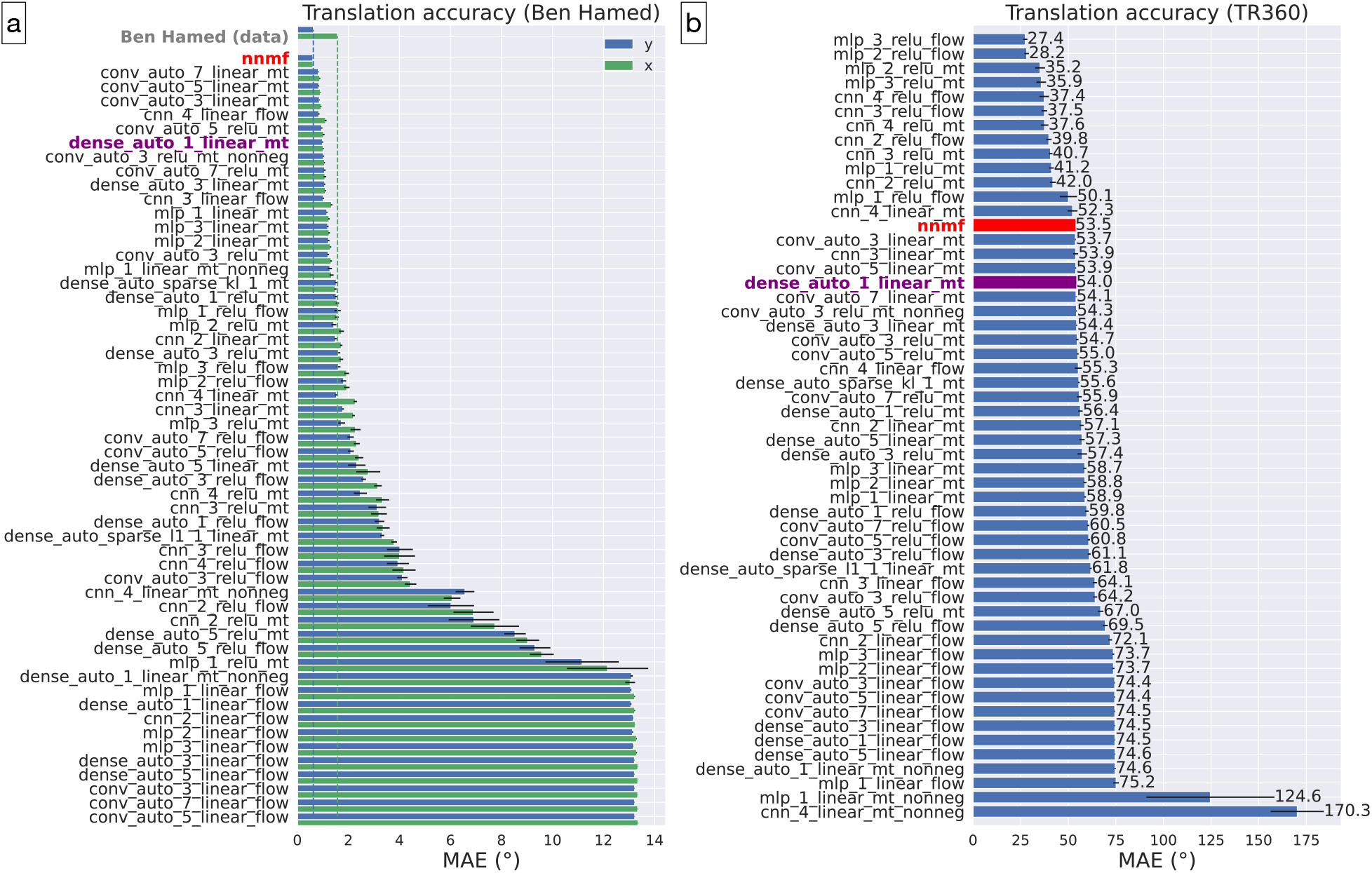
Self-motion estimation performance across models and datasets. **a**, Accuracy (MAE) of decoded translation direction from the hidden layer activations of ANNs and NNMF. Decoding was performed using 10-fold cross-validation on a translational dataset (30° heading eccentricity). Mean horizontal (*x*) and vertical (*y*) errors from MSTd neurons (Ben Hamed et al., 2003) are shown at top, with dashed lines indicating these biological decoding error levels. Models are ordered by accuracy. **b**, Performance on *T* + *R* test samples from the *T R*360 dataset. For accuracy-optimized models, values represent the error of the network output; for autoencoders and NNMF, values represent decoding error from the bottleneck layer. Error bars represent 95% CIs.

The T+R optic flow patterns used to train the ANNs are substantially more complex than the pure translation used in the Ben Hamed dataset. When we evaluated self-motion estimation on 3,015 TR360 test samples not used to fit the models, errors increase markedly overall (Figure 3b). Unsurprisingly, the 11 models achieving *<* 50° MAE are accuracy-optimized with nonlinear ReLU activation functions. This indicates that while linear models are capable of achieving MSTd-like decoding accuracy on simple translational patterns, they demonstrate poor generalization to more complex T+R flow. Finally, the mean decoding error of NNMF (53.7°) is comparable to several other autoencoder variants.

### Optic flow tuning metrics

Next, we characterized five key optic flow tuning properties of the model units and compared them to neurophysiological data. To provide an intuition for these properties before quantifying the correspondence across the entire 54-model population, we selected three exemplar models for qualitative comparison: a single-layer autoencoder (*dense_auto_1_linear_mt*), a deeper accuracy-optimized model (*cnn_3_relu_mt*), and NNMF. Figure 4a (top panel) shows the distribution of preferred translation directions for the population of MSTd neurons from Takahashi et al. (2007), with the corresponding preferences of units in the three exemplar models displayed underneath. Consistent with the neural data, autoencoder units exhibit a bimodal distribution with more units preferring lateral headings (left: 0/360°, right: 180°). Also consistent with the MSTd data, units in *dense_auto_1_linear_mt* demonstrate a bias toward forward (0–180°) compared to backward (180-360°) directions. NNMF exhibits qualitatively similar patterns, albeit with increased contrast between the peaks and troughs, indicating fewer neurons tuned to nearly straight ahead (90°) or backward (270°). As we showed previously, however, the units accuracy-optimized CNN demonstrate a substantial bias toward backward azimuths (Layton and Steinmetz, 2024; Layton et al., 2024).

**Figure 4.**
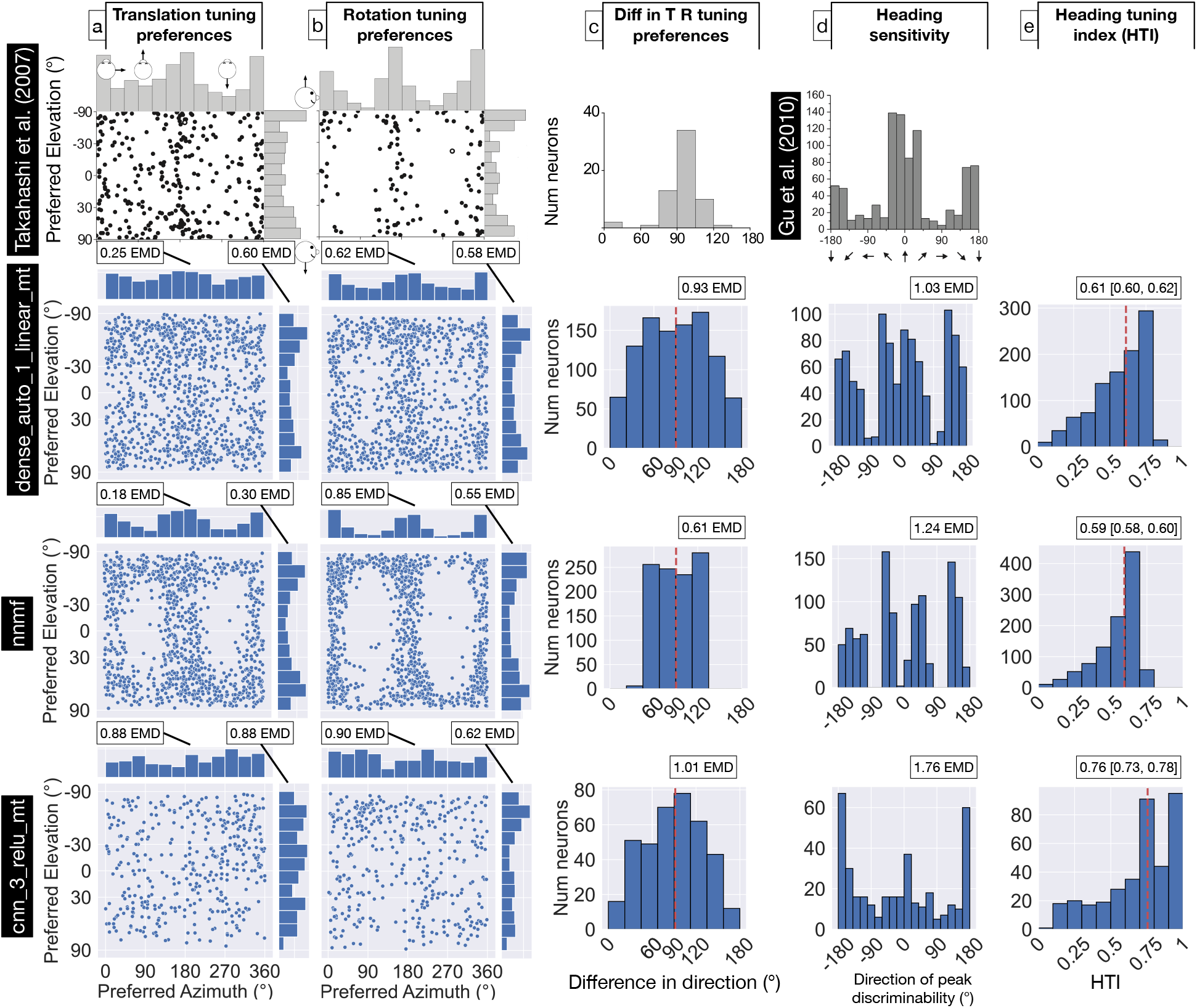
Optic flow tuning properties of MSTd neurons and exemplar model units. **a**, Preferred translation azimuth (top) and elevation (bottom) for a population of MSTd neurons (Takahashi et al., 2007) and units from three exemplar models: a linear autoencoder (*dense_auto_1_linear mt*), NNMF, and an accuracy-optimized CNN (*cnn_3_relu_mt*). Adjacent bar charts show marginal azimuth and elevation preferences. Labeled earth mover’s distances (EMD) indicate alignment between the annotated model distribution and the corresponding neural distribution (top panel). **b**, Rotation tuning preferences for the same populations. **c**, Difference between preferred translation and rotation azimuths for individual units. **d**, Distribution of headings yielding peak discriminability (maximum tuning slope) for MSTd (Gu et al., 2010) and model units. Dashed vertical lines show population medians. **e**, Heading Tuning Index (HTI) distributions.

To quantify the similarly between the MSTd and model distributions (*neural alignment*), we computed the earth mover’s distance (EMD; see Materials and Methods), where a value of 0 indicates an identical distribution and larger values indicate poorer correspondence. Consistent with visual inspection, NNMF achieves the lowest EMD (0.18), closely followed by the autoencoder (0.25). The accuracy-optimized CNN garners a considerably higher EMD (0.88). With respect to translation elevation, most MSTd neurons exhibit peak tuning toward vertical directions (±90°). Both the autoencoder and NNMF replicate this qualitative pattern, though NNMF achieves an improved match (lower EMD), whereas the accuracy-optimized CNN demonstrates a poor fit.

Figure 4b presents the rotation tuning preferences from Takahashi et al. (2007) alongside those of units in the three exemplar models. Similar to the neural data, azimuth rotational preferences in the autoencoder and NNMF exhibit increased polarization toward the cardinal axes compared to translation (see Figure 4-1 for percentages of neurons in each model that garner tuning nearby each cardinal axis). This effect is so pronounced in NNMF that the fit to the neural data degrades (EMD: 0.85); the more distributed preferences in the autoencoder provide a better fit (EMD: 0.62). Both the autoencoder and NNMF produce rotation elevation preferences comparable to the MSTd population. In contrast, rotation tuning in the accuracy-optimized CNN exhibits poor neural alignment for azimuth and elevation.

We considered how the difference in translation and rotation tuning preferences of each model unit compares to that of MSTd neurons from Takahashi et al. (2007) (Figure 4c). Most MSTd neurons exhibit a characteristic difference in preference of ≈ 90°, which the autoencoder (median: 90.28 [87.58, 92.97] CI), NNMF (median: 90.65 [90.27, 91.03]), and the CNN (median: 86.57 [77.98, 95.15] CI) all replicate. Of the three exemplar models, NNMF’s narrow distribution yields the closest neural alignment (EMD: 0.61), whereas the ANNs both produce much broader distributions.

We included a “heading sensitivity” metric in our analysis, which quantifies the translation azimuth at which a unit’s tuning curve yields its maximum slope (see Materials and Methods). The population of MSTd neurons recorded by Gu et al. (2010) yields a distribution of headings at which sensitivity is greatest (“direction of peak discriminability”) that peaks close to straight-ahead headings (0°; Figure 4d; top panel). A secondary peak arises around straight backward headings (±180°), and few units are most sensitive to lateral headings (±90°). Among the three exemplar models, the autoencoder achieves the best neural alignment (EMD: 1.03). While the distribution garnered by NNMF bears some similarity to that of the autoencoder, the neural alignment is weaker (EMD: 1.24) because fewer neurons possess peak heading sensitivity within 20° of the straight ahead (0°) and the population shows an increased lateral bias. The accuracy-optimized CNN yields a poor match due to its dominant backward bias (± 180°).

Finally, we considered the heading tuning index (HTI), which measures the strength of heading tuning and ranges from 0 to 1 (weak to strong tuning; see Materials and Methods). Figure 4e shows skew across the model populations toward stronger tuning. None of the model exemplars capture the mean HTI of 0.48 ± 0.16 SD found in MSTd (Gu et al., 2006), though NNMF (med: 0.59 [0.58, 0.60] 95% CI) and the autoencoder (med: 0.61 [0.60, 0.62] 95% CI) come closer than the CNN (med: 0.76 [0.73, 0.78] 95% CI). See Figure 4-2 for HTI values for every model.

### Neural alignment of optic flow tuning preferences across models

We sought to compare the neural alignment achieved by all 54 ANNs and NNMF with respect to the five optic flow tuning metrics. For each model, we report the mean and 95% confidence intervals (CIs) of the neural alignment (EMD) computed over 25 independent training repetitions. This sampling process facilitates comparison by quantifying the consistency in neural alignment while making explicit the variability caused by stochastic optimization and differing random choice of initial ANN weights.

Figure 5 shows the neural alignment of each model with respect to translation and rotation tuning preference (see Figure 5-1, Figure 5-2, and Figure 5-3 for other tuning metrics). With respect to translation azimuth, NNMF achieves the closest neural alignment to MSTd (mean EMD: 0.18 [0.13, 0.23]), following by two shallow, linear autoencoders *dense_auto_1_linear_mt* (mean EMD: 0.25 [0.14, 0.36]) and *dense_auto_-sparse_l1_1_linear_mt* (mean EMD: 0.25 [0.11, 0.39])(Figure 5a). The top five models are shallow, linear architectures that implement either the accuracy-optimized and motion-reconstruction objective. Broadening the focus to top ten most neurally aligned models, all but one process MT-encoded input signals rather than raw optic flow.

**Figure 5.**
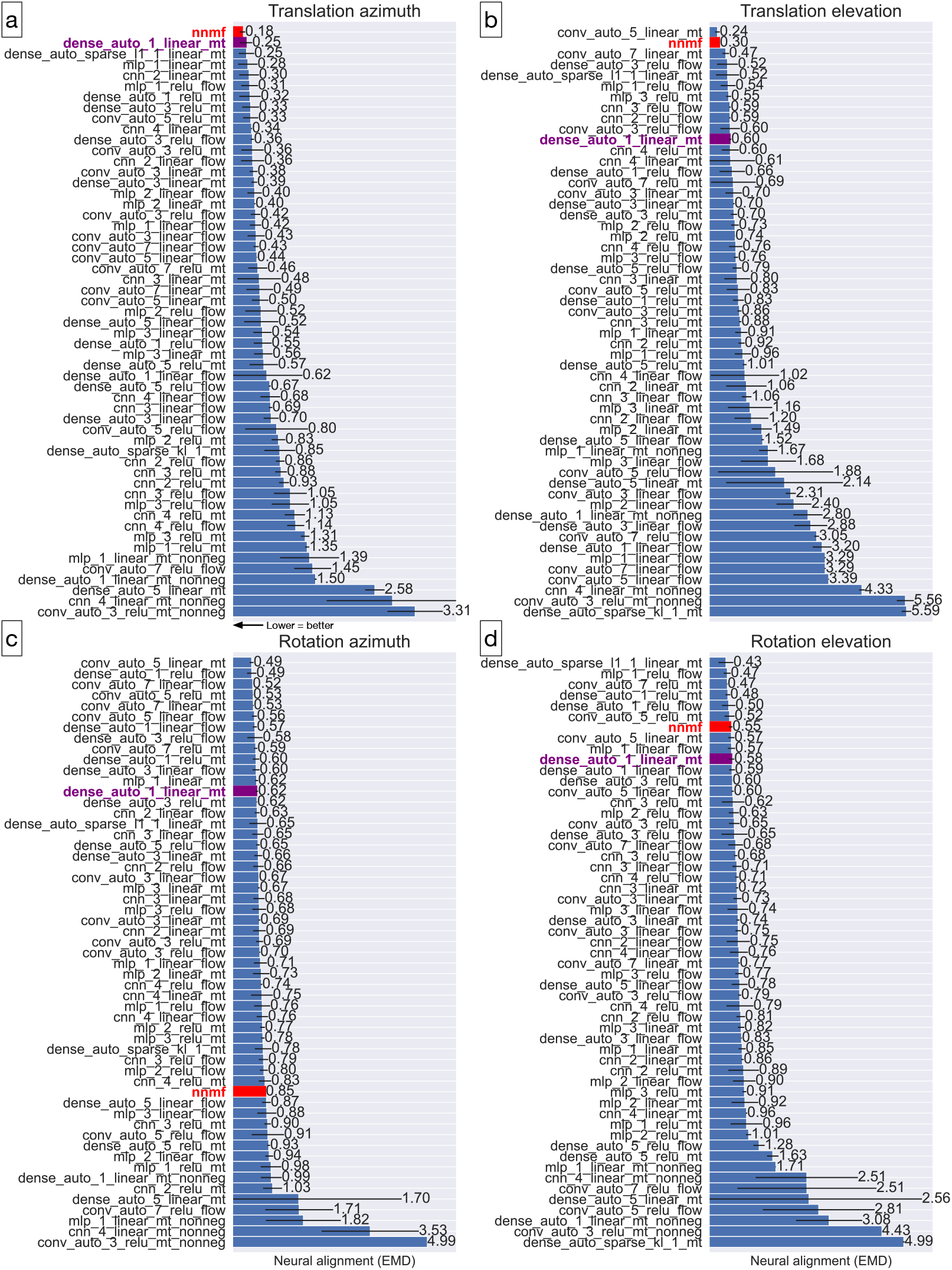
Neural alignment (EMD) of optic flow tuning preferences across 54 ANNs and NNMF. **a**, Alignment between model and neural distributions for translation azimuth. Models are ordered by mean alignment score. **b**, Alignment for translation elevation. **c**, Alignment for rotation azimuth. **d**, Alignment for rotation elevation. In all panels, lower EMD values represent closer correspondence to MSTd data. Error bars indicate 95% CIs across 25 training repetitions.

For translation elevation preference (Figure 5b), a convolutional autoencoder garners the closest neural alignment (mean EMD: 0.24 [0.18, 0.31]), followed by NNMF (mean EMD: 0.30 [0.29, 0.31]). Among the top five models, all optimize the motion-reconstruction objective, and all but one are linear and process MT-encoded input. When considering the top ten models, however, the computational attributes are more heterogeneous.

With respect to rotation azimuth preference, the top ten models are all autoencoders (Figure 5c). Interestingly, NNMF yields poor neural alignment compared to most ANNs. This is due to the strong clustering in preferences around lateral headings (Figure 4b). Finally, a shallow sparse autoencoder (*dense_auto_sparse_- l1_1_linear_mt*) produces the closest match to MSTd rotation elevation preference (Figure 4d). Among the ten most neurally aligned models for this metric, eight implemented the motion reconstruction objective and seven process MT-encoded input.

Along these four tuning preference dimensions, the ANNs with the non-negative weight constraint consistently yield the poorest neural alignment, as does the sparse autoencoder with the KL-divergence penalty (*dense_auto_sparse_kl_1_linear_mt*).

Before moving on, it is noteworthy that we explored whether there is a relationship between self-motion decoding accuracy and neural alignment, but found no such association (*R*^2^ = 0.02; Figure 5-1).

### Relative neural alignment across models and metrics

We examined the overall neural alignment of each model across all five tuning metrics. For each model and metric, we established the neural alignment relative to a baseline model and calculated the geometric mean of those relative EMD values. This yields a single relative neural alignment score per model (“mean relative alignment score”). We repeated this process for the 25 repetitions of each model to determine the mean and 95% CI. As a baseline, we selected the shallowest, linear, dense autoencoder (*dense_auto_1_linear_mt*), which consistently achieves close neural alignment across the five metrics.

Figure 6a shows the relative alignment score for each model. By construction, the baseline model achieves a score of 1.0; models with relatively closer (better) neural alignment garner scores less than 1.0, while high scores indicate poorer correspondence. Only NNMF (mean score: 0.78 [0.60, 1.02]) and the autoencoder with a L1 sparsity constraint (*dense_auto_sparse_l1_1_linear_mt*; mean score: 0.90 [0.63, 1.3]) produce lower mean relative alignment scores than the baseline. Notably, the top ten ANNs are all autoencoders that process MT-encoded input. While the top three models are linear with dense connectivity, nonlinear and convolutional models also produce competitive scores within the top ten. Consistent with translation and rotation preference alignment (Figure 5), models with the non-negative weight constraint yield poor relative overall alignment.

**Figure 6.**
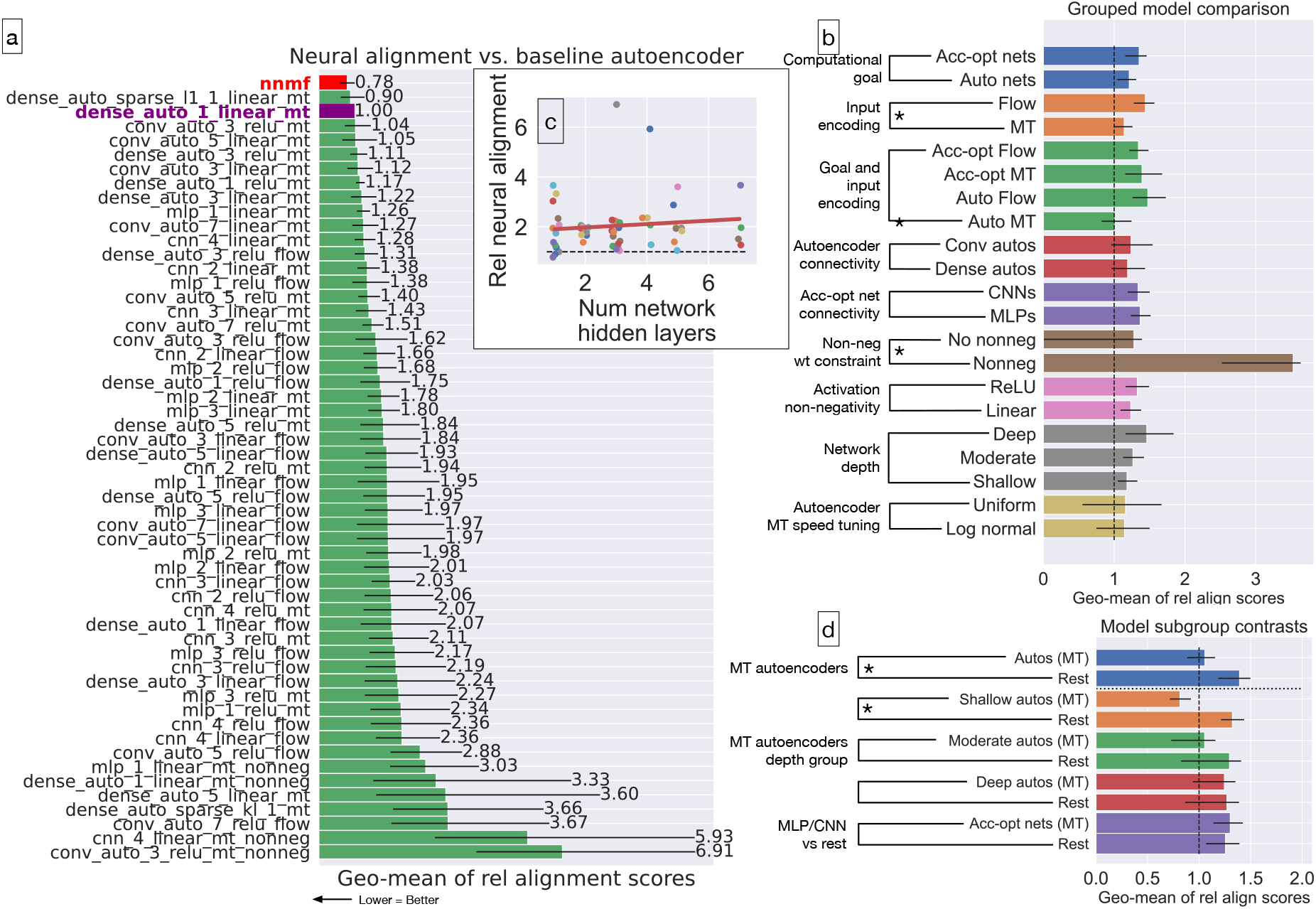
Relative neural alignment scores across the model population. **a**, Relative alignment score for each of the 54 ANNs and NNMF. Scores represent the geometric mean of the EMD values across all five tuning metrics, normalized to the baseline autoencoder (*dense_auto_1_linear_mt*, score = 1.0). Lower scores indicate closer alignment to MSTd data. **b**, Mean relative alignment scores of model subgroups categorized by computational attributes. **c**, Relative alignment score as a function of the number of hidden layers in the ANNs. The dashed line represents a linear regression (*R*^2^ = 0.01). **d**, Comparison of relative alignment scores for MT-encoded autoencoders (grouped by depth) against the remaining ANN population. Error bars show 95% CI across 25 training repetitions.

To determine the computational attributes that significantly impact neural alignment, we computed the mean relative alignment scores within different model subgroups (Figure 6b). We found that models processing MT input generate significantly closer neural alignment compared to those processing raw optic flow (MT: 1.5 [1.32, 1.69], flow: 2.0 [1.83, 2.17]). Moreover, models without the non-negative weight constraint yield significantly closer neural alignment than those restricted by the constraint (no constraint: 1.73 [1.58, 1.89], non-negative constraint: 4.51 [3.02, 6.74]). Across the ANNs, the use of linear versus nonlinear activations does not affect relative alignment (t(46.98)=−1.22, p=0.23). Similarly, neither the computational objective (t(36.08)=−1.65, p=0.11) nor connectivity—within either the autoencoders (t(22.84)=−0.27, p=0.79) or accuracy-optimized networks (t(21.65)=−0.10, p=0.92)—significantly impacts alignment. Relative neural alignment is not associated with the number of hidden layers in the ANNs (Figure 6c; *R*^2^ = 0.01). It is notable that relative neural alignment is also not associated with the accuracy of self-motion estimates (Figure 6-1).

We probed the MT motion signal advantage further to determine whether the way speed tuning is modeled impacts neural alignment. To that end, we trained a variant of the dense autoencoder ANNs that possess uniform rather than the log-normal MT speed tuning used heretofore (Nover et al., 2005). This change has no effect on the relative neural alignment (Figure 6b; t(9.76)=−0.04, p=0.97).

Taken together, Figure 6a–b suggest a neural alignment advantage of autoencoders that process MT-encoded input signals. We examined this more closely by comparing the relative alignment scores of this specific group compared to the other ANNs. This group of autoencoders yields significantly lower (better) relative neural alignment scores than the other ANNs (Figure 6d). In particular, both the shallow and medium depth, but not deep, autoencoders garner significantly lower relative alignment scores. Crucially, accuracy-optimized MLPs and CNNs that process the MT signals do not exhibit a significant difference in alignment compared to the other models. This indicates that the significantly better alignment achieved by the ANNs with MT rather than raw optic flow inputs stems from the autoencoders (Figure 6b).

In sum, the relative alignment scores reveal a clear hierarchy in biological consistency: ANNs that combine a MT-like motion representation with a reconstruction-based objective achieve the highest alignment with MSTd, whereas accuracy-optimized ANNs and those with explicit non-negativity constraints diverge from MSTd optic flow tuning properties.

### Dimensionality reduction and sparsity

Beyeler et al. (2019) hypothesize that dimensionality reduction and sparsity may represent key computational principles that underlie optic flow tuning properties in MSTd. We used our modeling framework to investigate this proposal. Given that autoencoders offer explicit control over the extent of the dimensionality reduction performed on the input, we conducted an experiment using the baseline autoencoder (*dense_ - auto_1_linear_mt*) wherein we systematically varied the dimensionality by changing the number of units in the hidden layer. Figure 7a shows the neural alignment (EMD) for the translation and rotation preference metrics as a function of the autoencoder dimensionality. Contrary to the hypothesis, the neural alignment improves rather than degrades as the learned dimensionality increases. Interestingly, the alignment plateaus after a dimensionality of 32–256, depending on the metric. Notably, alignment when the learned dimensionality matches that of the MT input signal (9,000 units) is largely unchanged compared to that achieved with autoencoders with much lower dimensionality.

**Figure 7.**
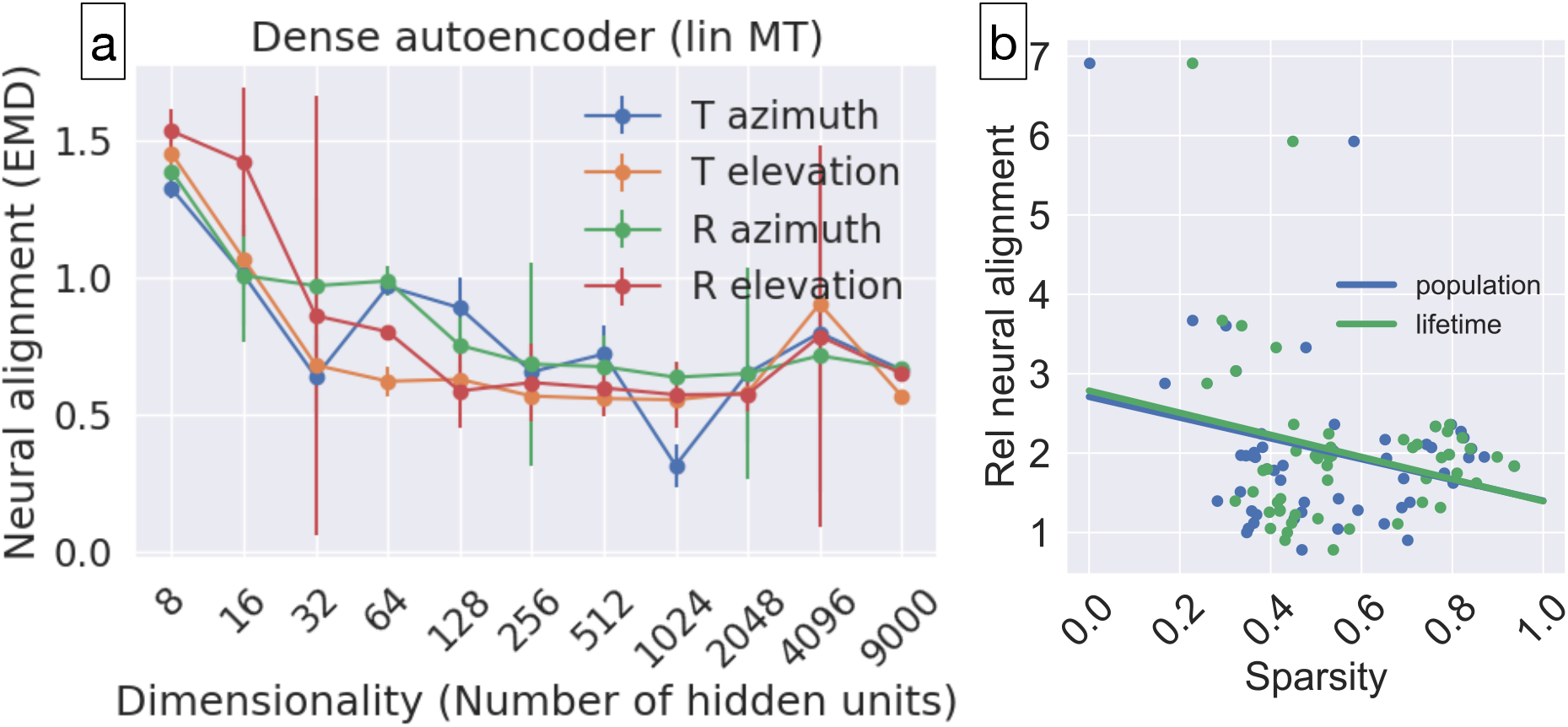
Effect of dimensionality and sparsity on neural alignment. **a**, Neural alignment (EMD) for translation and rotation azimuth preferences as a function of the number of hidden units in the baseline autoencoder (*dense_auto_1_linear_mt*). Error bars represent 95% CIs. **b**, Relationship between relative alignment score and two sparsity metrics: population sparsity (blue) and lifetime sparsity (green) across all 54 ANNs. Each data point represents the mean score for a specific model.

Additionally, we quantified the sparsity in the 54 ANNs using the population and lifetime sparsity metrics (see Materials and Methods and Figure 7-1). Both metrics range from 0 to 1, where values closer to 1 indicate a sparse code and values closer to 0 indicate a dense code. For the population metric, a sparse code means that few neurons activate to a typical optic flow sample; for the lifetime metric, it means that a typical neuron is activated by only a few optic flow samples in the dataset. Figure 7b shows both the sparsity and relative alignment score of each ANN computed on the TR360 dataset. Linear regressions revealed that neither the population (*R*^2^ = 0.07) nor lifetime (*R*^2^ = 0.06) sparsity is associated with neural alignment.

Our analysis suggests that dimensionality reduction and sparsity alone may not drive alignment with optic flow tuning in MSTd. Instead, the computational objective coupled with the MT input encoding may serve as superordinate organizing principles.

## Discussion

Our analysis of 54 ANN architectures demonstrates that computational objective and input encoding are primary drivers of neural alignment with MSTd optic flow tuning properties. While the choice of computational objective alone did not significantly affect overall alignment among the models simulated (accuracy optimization vs. input reconstruction), the inclusion of MT-like speed and direction tuned inputs consistently improved model correspondence to MSTd data—an effect driven primarily by the autoencoder ANNs (Figure 6b). Among the ANNs tested, we found that autoencoders tasked with reconstructing MT-like motion signals yield the highest alignment with MSTd optic flow tuning properties. The NNMF model (Beyeler et al., 2016), which incorporates both components, achieved comparable overall alignment to the MT autoencoders, suggesting that these two factors may underlie why NNMF effectively captures MSTd tuning properties. Interestingly, the difference in mean neural alignment with autoencoders that process optic flow directly suggests that these autoencoders do not inherently develop MT-like representations. Other constraints inherited from NNMF—including non-negativity, dimensionality reduction, and sparsity—offer no additional improvement in the ANNs. Taken together, these findings suggest that a reconstruction objective paired with a MT-like input encoding promotes MSTd-like optic flow tuning; the additional mechanisms of the NNMF model may not be essential.

### Comparison to existing models

Our results offer a parsimonious alternative to recent ANN models of optic flow processing in the dorsal stream. For instance, Vafaii et al. (2024) demonstrated that a hierarchical Variational Autoencoder (VAE) trained on optic flow patterns captures a large proportion of the variance of MT neural responses (R≈0.5; Cui et al., 2013). Their VAE includes 21 “latent groups” or effective layers to process motion signals at multiple spatial scales—an architecture that is substantially deeper than the autoencoder variants simulated here (maximum: 8 layers). This depth contrasts with our finding that the mean neural alignment with MSTd is highest in the shallowest architectures, with deeper networks showing a trend toward reduced alignment (Figure 6d). While it is possible that the stochastic, sampling-based nature of the VAE requires greater depth than our deterministic, weighted-sum framework, a more likely explanation involves the input representation. We provide our models with optic flow already transformed by a mechanistic MT model, whereas Vafaii et al. (2024) trained their VAE directly on optic flow vectors. Therefore, some of the depth in the VAE may be dedicated to learning the MT-like features that are provided a priori to our MSTd models. This suggests that while modeling the entire hierarchy from raw optic flow to MSTd may benefit from depth, the transformation from MT to MSTd properties is efficiently captured by shallow, reconstruction-driven architectures. The authors do not simulate VAE variants with different numbers of layers, so the necessity of the 21 layers for modeling the MT data of Cui et al. (2013) remains unclear.

Our results also contrast with DorsalNet, a 3D CNN optimized for self-motion estimation (Mineault et al., 2021). Although a direct comparison is challenging because Mineault et al. focused on correlations with neural recordings rather than tuning properties, several key distinctions are noteworthy. First, DorsalNet utilizes a task-optimization objective similar to our accuracy-optimized ANNs. Consistent with our finding that reconstruction objectives improve neural alignment, Vafaii et al. (2024) reported that their VAE captured approximately double the variance in the Cui et al. (2013) MT dataset compared to DorsalNet. Second, DorsalNet operates directly on raw image pixels, whereas the models discussed here—including the NNMF model (Beyeler et al., 2016), the VAE (Vafaii et al., 2024), and our autoencoder variants—utilize optic flow as an input representation. Whether computational consequences stem from these differing front end input representation warrants investigation. Finally, DorsalNet consists of 14 layers, exceeding the 8-layer maximum in our study. This increased depth is likely a functional requirement for learning early- and mid-level motion features from raw pixels, whereas our use of MT-like input encoding allows for a shallower, more computationally efficient transformation into MSTd-like representations.

### Computational goals and algorithmic constraints of the dorsal stream

The reconstruction objective of the autoencoder ANNs offers a biologically plausible framework for unsupervised learning, as it requires only local access to the feedforward signals from area MT (Van Essen and Maunsell, 1983). By relying on the input signal itself, this strategy bypasses the need for explicit self-motion ground truth labels. This approach aligns with a growing body of unsupervised models (Zhuang et al., 2021; Nayebi et al., 2021; Konkle and Alvarez, 2022; Vafaii et al., 2024) and predictive coding (Clark, 2015; Millidge et al., 2021).

Consistent with Marr (1982)’s hierarchy, our findings characterize the computational *goal* of MSTd at an algorithmic level. We probed neural alignment of autoencoders using the bottleneck layer, halfway between the encoder and decoder subnetworks. Here, the decoder primarily provides the reconstruction error to backpropagate and refine the bottleneck representations. In the cortical dorsal stream, such feedback could be implemented via local learning rules (Whittington and Bogacz, 2017; Bienenstock et al., 1982; Oja, 1982; Chen et al., 2022) or feedback signals (Lillicrap et al., 2016), removing the need for a distinct decoder.

### Poor accuracy of self-motion estimation in autoencoders

Substantial evidence indicates that MSTd is involved in self-motion perception (Britten and Van Wezel, 1998; Gu et al., 2012). The direction of translation can be linearly decoded from MSTd populations with an accuracy of ≈1° (Ben Hamed et al., 2003), matching the precision of human heading judgments (Warren et al., 1988). While the ANNs reproduce this high level of precision when tested on the stimuli of Ben Hamed et al. (2003), they exhibit significantly higher mean error (≈50°) on the TR360 dataset (Fig. 3b).

This large discrepancy is likely attributable to the dramatic increase in task difficulty. Whereas the Ben Hamed stimuli restrict simulated self-motion to pure translation at a fixed 30° eccentricity, the TR360 dataset includes all possible headings across the unit sphere, frequently coupled with high-velocity rotations (up to 10°/s). Human performance is known to degrade significantly outside of foveal headings—reaching errors of 10–15° at the periphery (Gu et al., 2010; Cuturi and Macneilage, 2013)—and these errors further compound in the presence of simulated rotation (Royden et al., 1994). Given that TR360 includes a much broader range of optic flow patterns than stimuli used in existing psychophysical and neurophysiological studies, it is plausible that human heading judgments would demonstrate similarly elevated error rates under these challenging conditions.

While our models are limited to visual inputs in the present study, MSTd is a multisensory area that integrates vestibular signals (Gu et al., 2010; Bremmer et al., 1999). These non-visual signals offer an independent reference frame that may improve the accuracy of self-motion perception along peripheral headings and in the presence of rotation. Within an autoencoder framework, multi-sensory integration of visual and vestibular signals may contribute a form of ‘weak supervision,’ potentially reducing perceptual error under complex self-motion conditions.

### Sparsity and Equifinality

The lack of a systematic relationship between sparsity and neural alignment is particularly striking given the established role of sparse coding in models of the early visual system (Olshausen and Field, 1996; Bell and Sejnowski, 1997). In V1, sparsity is a primary driver of the emergence of localized, oriented receptive fields; however, our results suggest that for the more complex, integrative task of optic flow processing in MSTd, sparsity may represent an epiphenomenon rather than a functional mechanism. This is supported by our observation that explicitly enforcing sparsity in the autoencoder objective, either through L1 or KL-divergence penalties, did not yield an improvement over baseline models. Furthermore, our previous work (Layton et al., 2024) demonstrated that extreme sparsity, resulting from ‘dead neurons’ in ReLU-based accuracy-optimized networks, failed to produce MSTd-like tuning despite high task performance. Our most aligned models tend to exhibit moderate sparsity levels (≈0.5), comparable to the original NNMF model (Beyeler et al., 2016). Notably, Vafaii et al. (2024) assessed sparsity levels in their VAE as a function of the β coefficient, which scales the KL-divergence term. While the authors emphasize the sparse nature of these activations, the median sparsity across *β* values remains moderate (≈0.5; see their Figure 8c), similar to the NNMF model and the most aligned ANNs in the present study. This commonality across disparate modeling frameworks suggests that a moderate activation density may be a natural property of optic flow representations, further supporting our view that the enforcement of sparsity may not be a primary driver of MSTd-like tuning.

The emergence of identical functional properties from different modeling conditions is known as ‘equifinality’ (Carozza et al., 2023; Lenc and Vedaldi, 2015). Throughout our analysis, we repeatedly observed models producing convergent MSTd-like properties despite having considerably different mechanisms, such as varied activation functions, dense vs. convolutional connectivity, and differing speed-tuning distributions in the MT stage (Figure 6b). Additionally, almost all models exhibited a 90° difference in translation and rotation tuning preferences (Figure 4c). This suggests the pattern may be a mathematical inevitability of processing visual motion in a computationally efficient manner. Future neurophysiological studies that sample a broader range of self-motion conditions may help clarify whether such properties are truly equifinal or are artifacts of narrowly sampling the task space.

## Conclusion

In this study, we investigated the effectiveness with which different ANN architectures, constraints, and computational objectives lead to the emergence of MSTd-like optic flow tuning. Our results demonstrate that these tuning properties are most effectively captured by shallow autoencoders optimized to reconstruct motion signals from MT-like inputs. Whereas accuracy-optimized ANNs currently represent the leading models of the ventral stream, our findings suggest that the dorsal stream may prioritize an alternative computational objective focused on signal reconstruction.

## Acknowledgments

Sandia National Laboratories is a multi-mission laboratory managed and operated by National Technology & Engineering Solutions of Sandia, LLC (NTESS), a wholly owned subsidiary of Honeywell International Inc., for the U.S. Department of Energy’s National Nuclear Security Administration (DOE/NNSA) under contract DE-NA0003525. This written work is authored by an employee of NTESS. The employee, not NTESS, owns the right, title and interest in and to the written work and is responsible for its contents. Any subjective views or opinions that might be expressed in the written work do not necessarily represent the views of the U.S. Government. The publisher acknowledges that the U.S. Government retains a non-exclusive, paid-up, irrevocable, world-wide license to publish or reproduce the published form of this written work or allow others to do so, for U.S. Government purposes. The DOE will provide public access to results of federally sponsored research in accordance with the DOE Public Access Plan.

## 1 Supplementary Material

**Figure 4-1.**
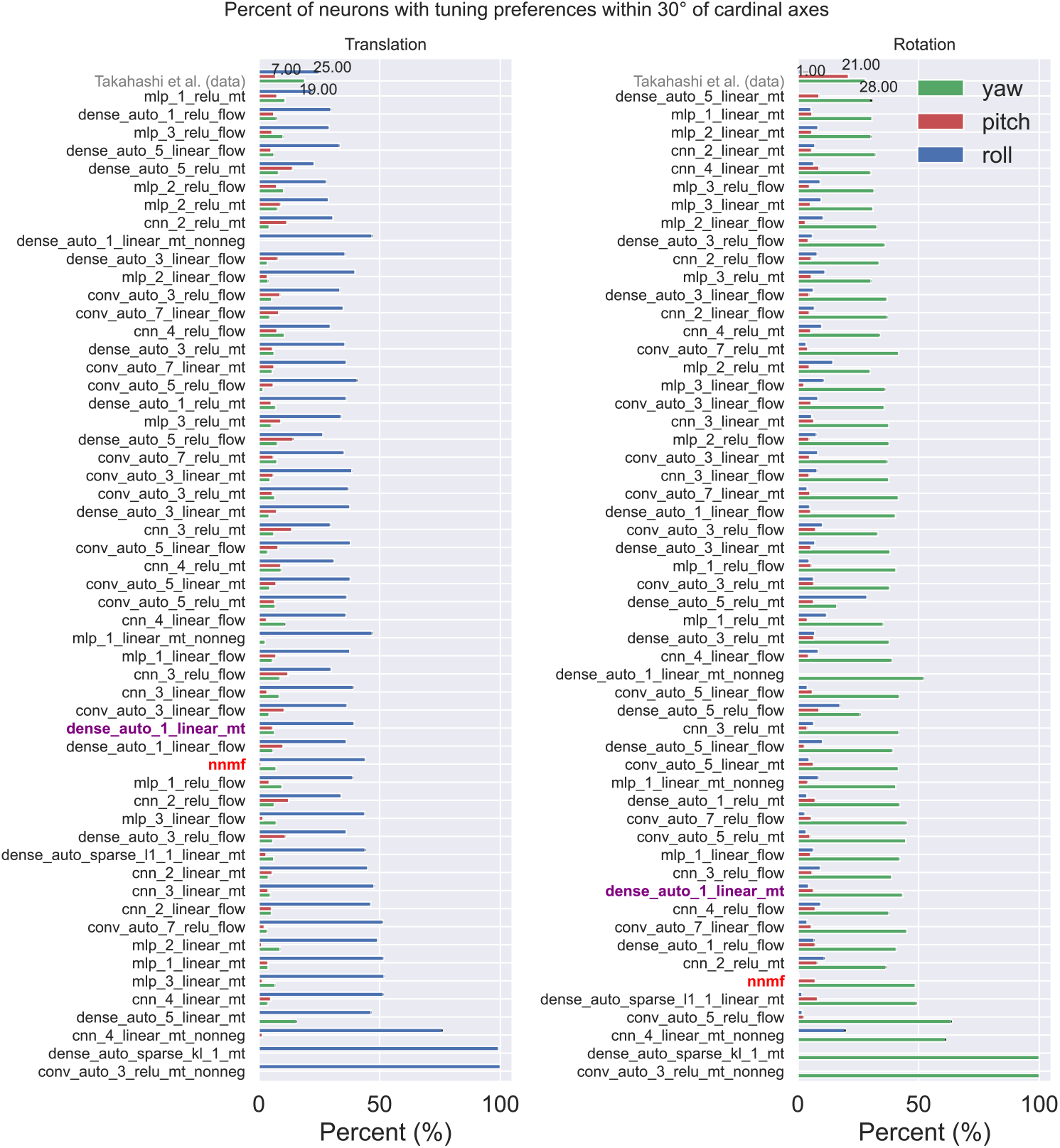
Preferred tuning alignment with cardinal axes. Percent of neurons in each model with tuning preferences within 30° of the cardinal axes for translation (left) and rotation (right). Gray bars indicate percentages from the MSTd population recorded by Takahashi et al. (2007). (Left) Vertical, forward-backward, and lateral axes are shown in blue, red, and green, respectively. (Right) Roll, pitch, and yaw axes are shown in blue, red, and green, respectively. Regardless of architecture, every model demonstrates a disproportionate preference for the vertical translation and yaw rotation axes.

**Figure 4-2.**
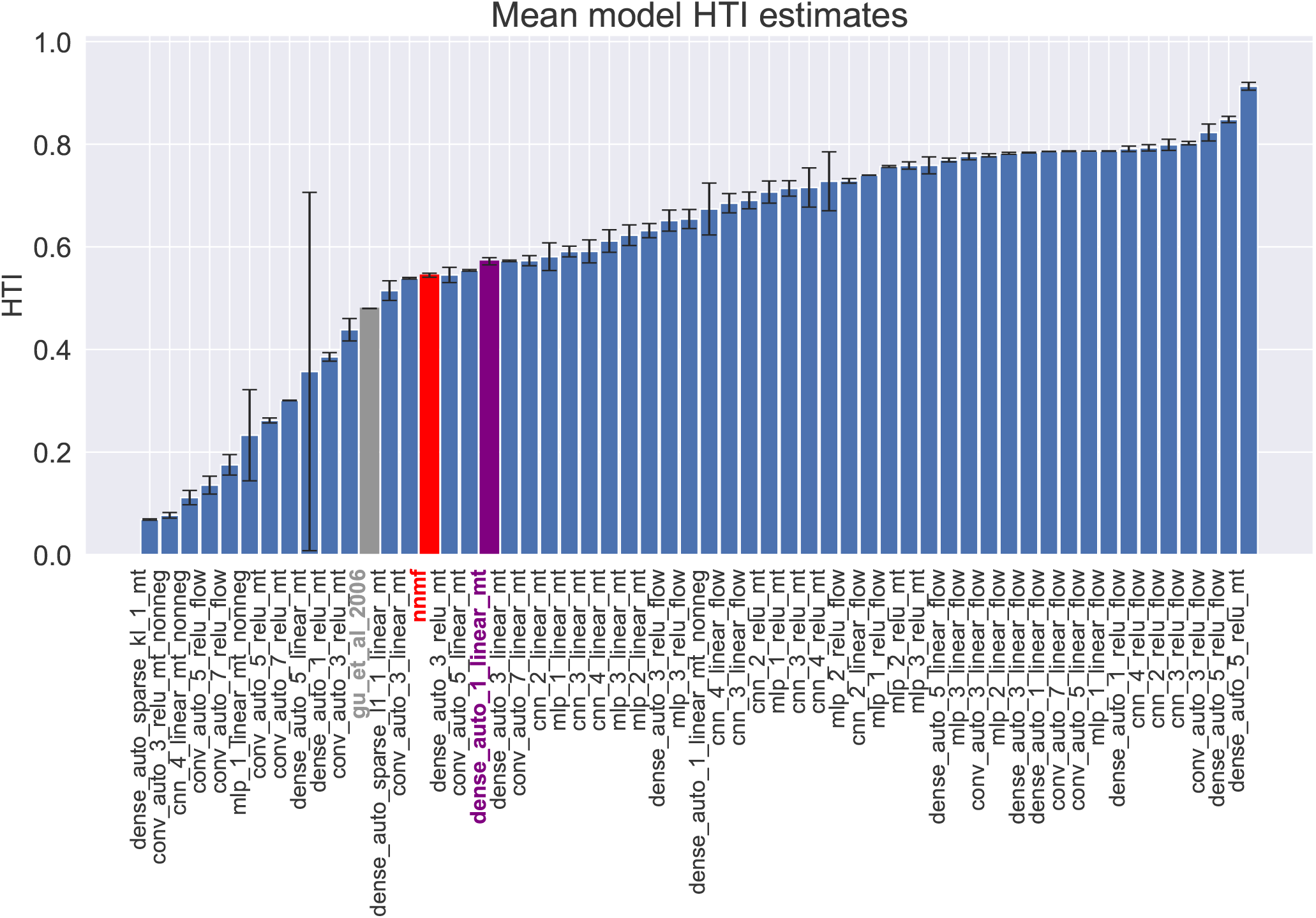
The mean heading tuning index (HTI) obtained on the TR360 dataset for each simulated model. Error bars show 95% CI across 25 training repetitions.

**Figure 5-1.**
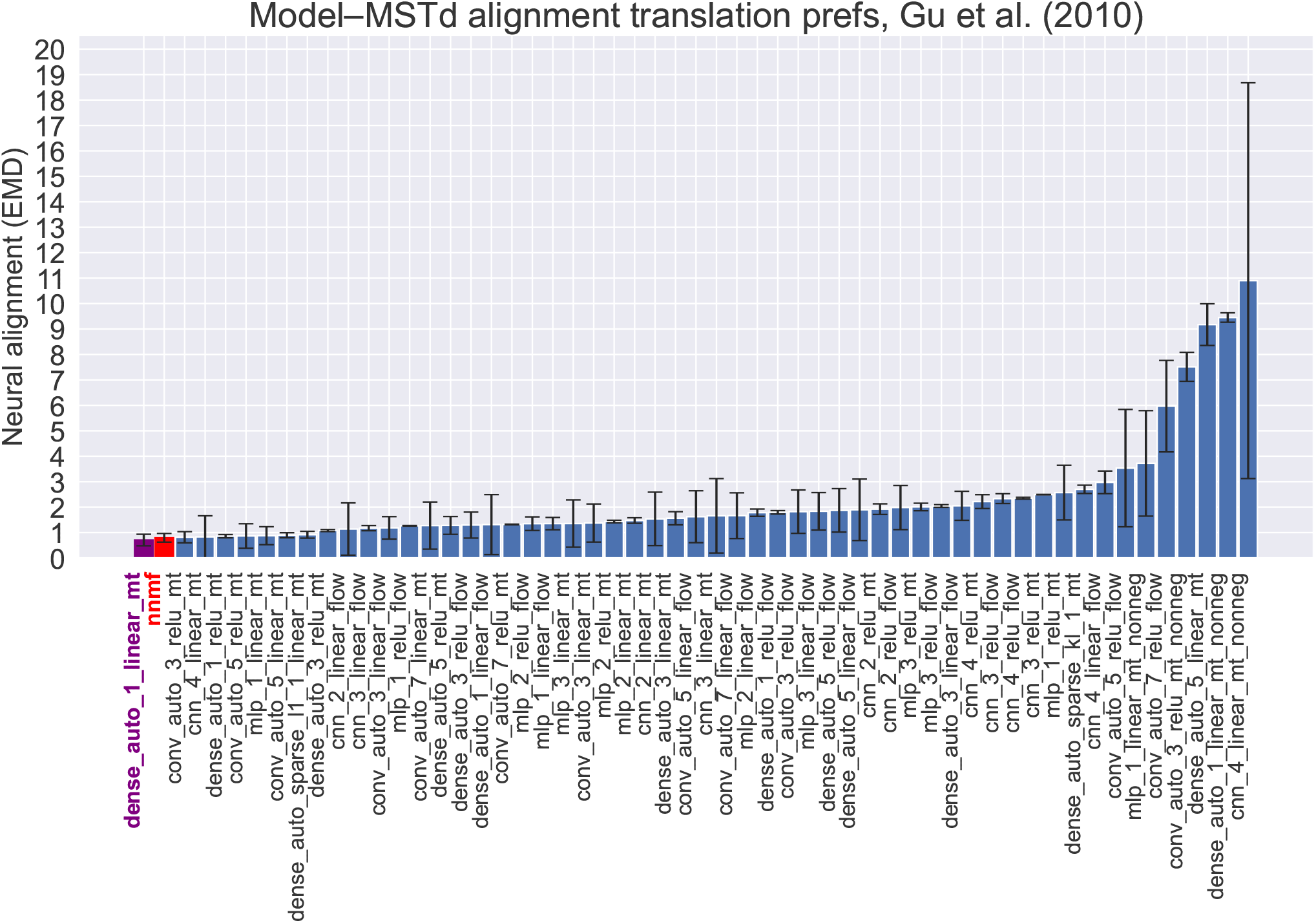
Alignment of model and neural translation azimuth distributions. Correspondence between simulated models and MSTd population data from Gu et al. (2010) for translation azimuth. Models are ordered by mean alignment score. Lower Earth Mover’s Distance (EMD) values represent closer distributional correspondence to the MSTd recordings. Error bars indicate 95% CIs across 25 training repetitions. Note that autoencoder architectures with MT-like input encoding consistently achieve the lowest EMD scores, indicating the highest alignment with physiological translation preferences.

**Figure 5-2.**
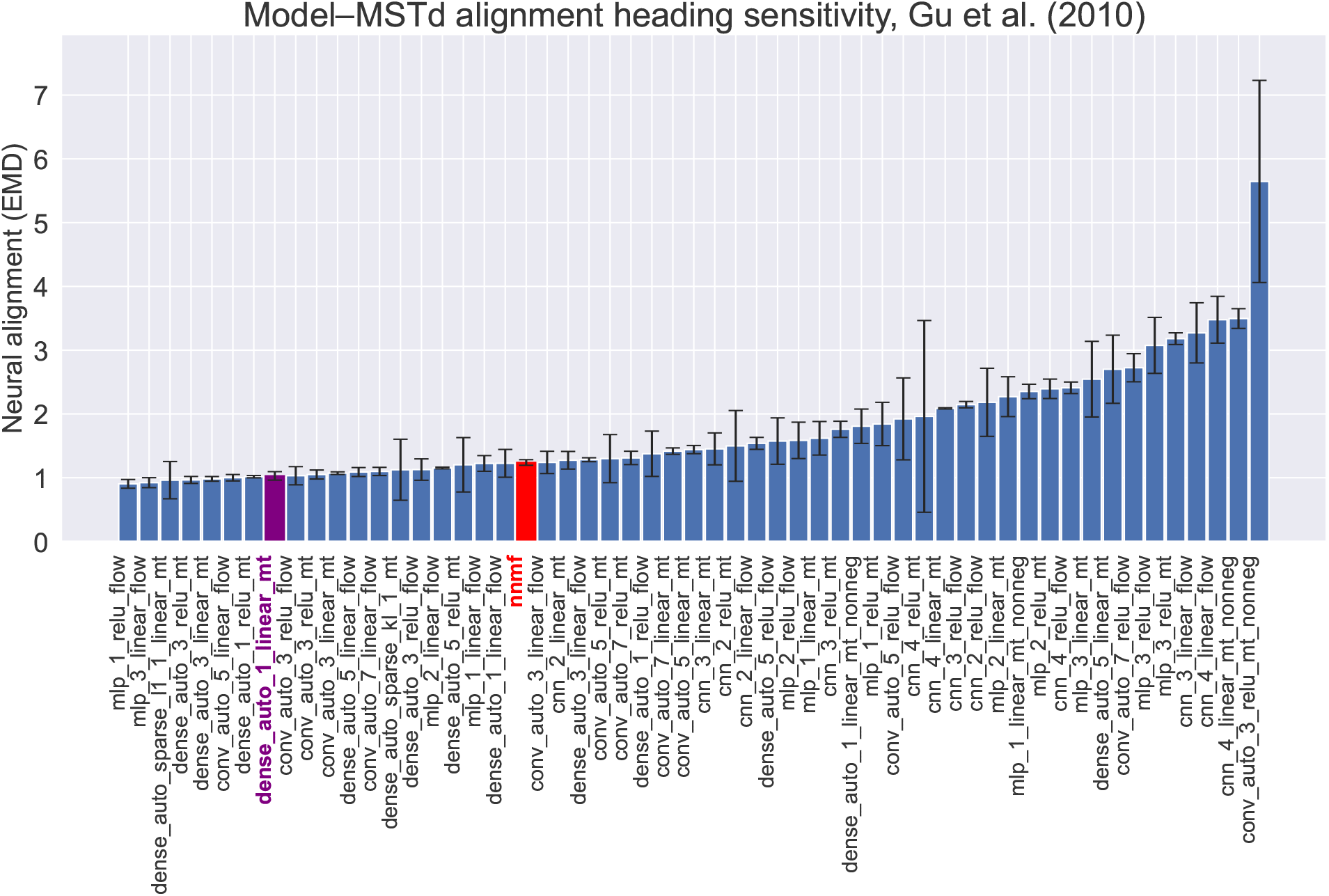
Alignment of model and neural heading sensitivity distributions. Correspondence between simulated models and MSTd population data from Gu et al. (2010). Models are ordered by mean alignment score. Lower EMD values represent closer distributional correspondence to the MSTd recordings. Error bars indicate 95% CIs across 25 training repetitions.

**Figure 5-3.**
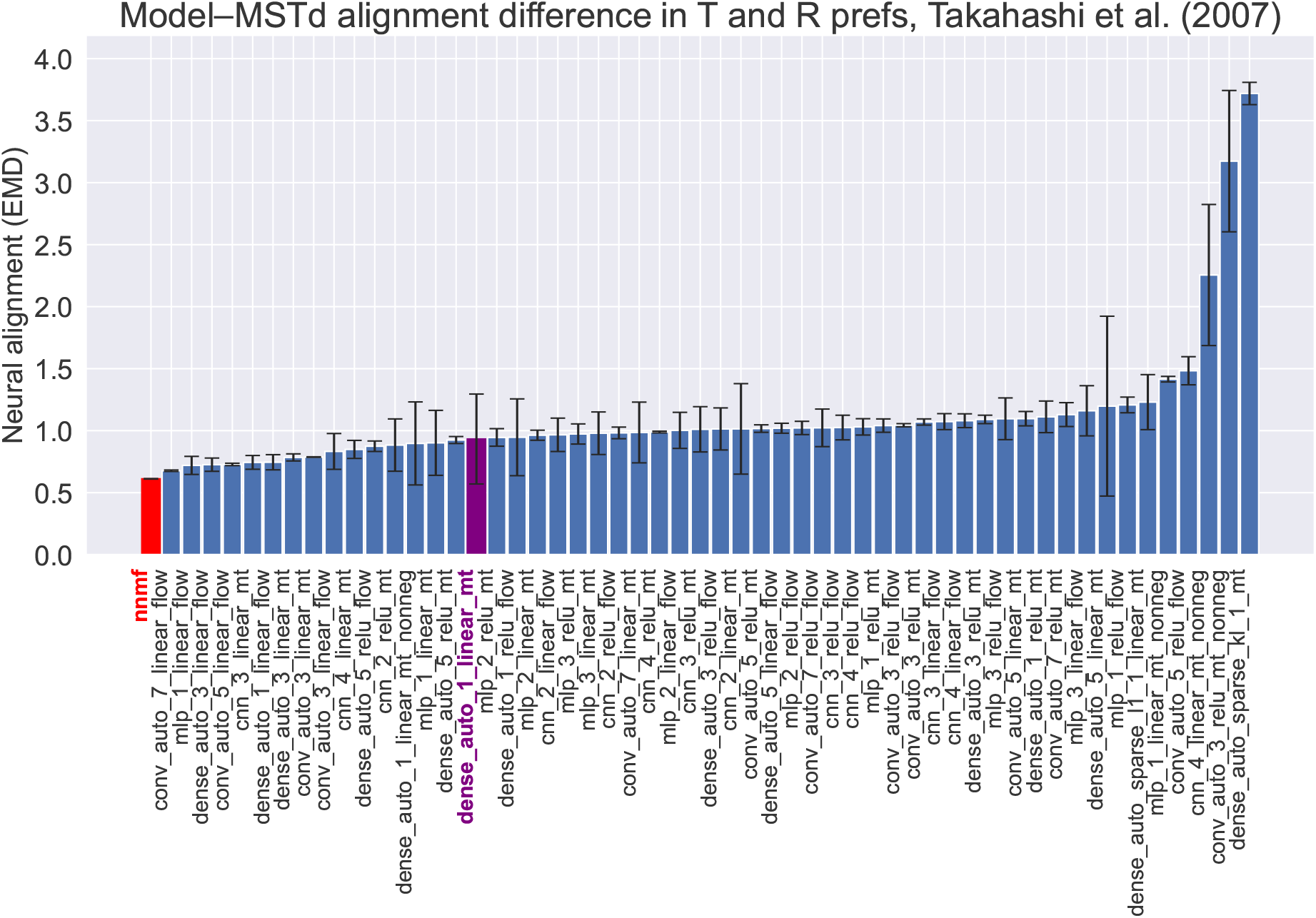
Alignment of translation-rotation preference differences. Correspondence between simulated models and MSTd population data from Takahashi et al. (2007) regarding the angular difference between preferred translation and rotation directions. Models are ordered by mean alignment score, where lower EMD values represent closer distributional correspondence to the physiological recordings. Error bars indicate 95% CIs across 25 training repetitions. Note that the majority of models achieve a low EMD (near 1.0), reflecting a distribution centered around the ≈90° average difference characteristic of MSTd neurons, regardless of architectural constraints.

**Figure 6-1.**
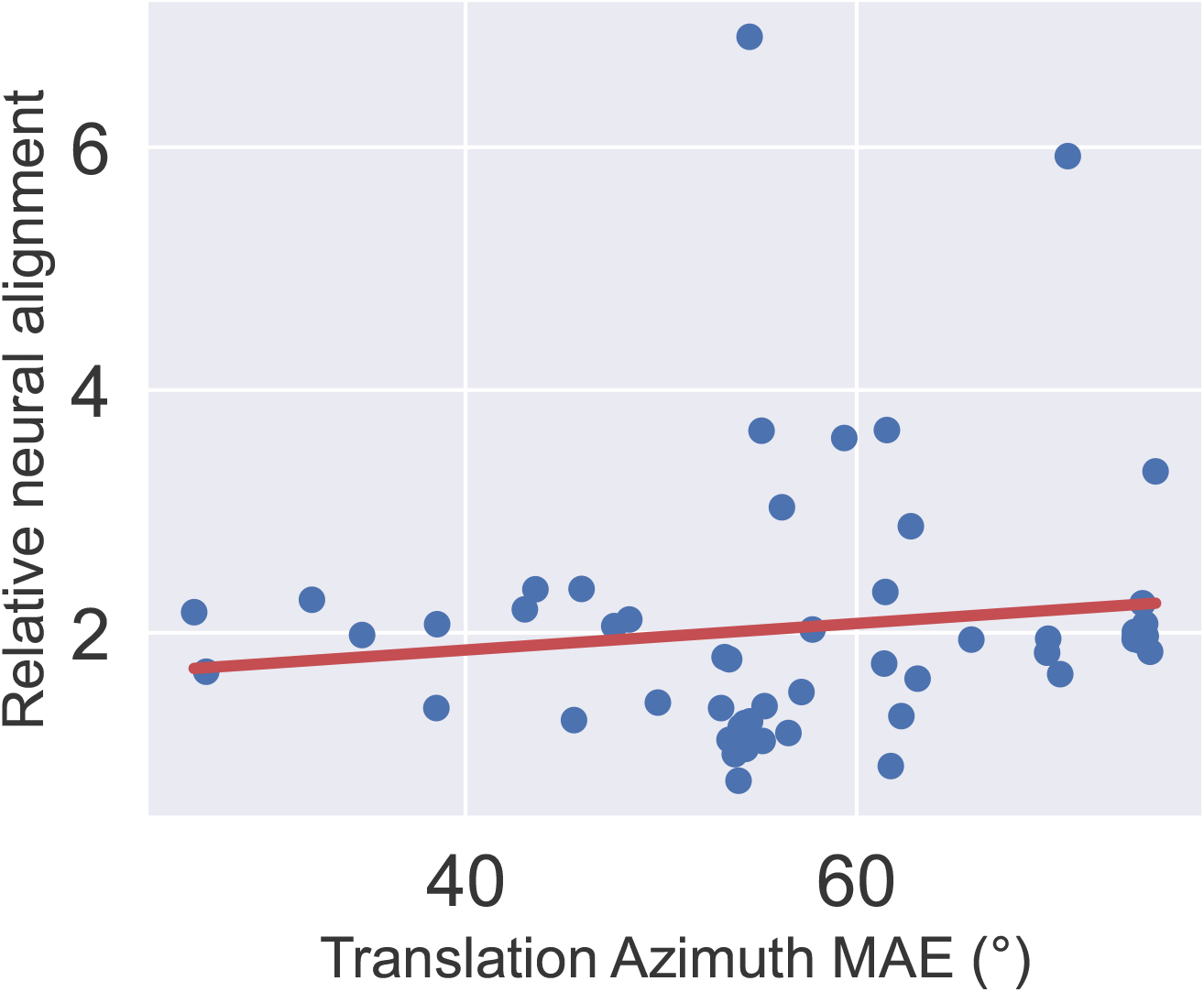
Relationship between self-motion accuracy and neural alignment. Mean Absolute Error (MAE) of model translation azimuths plotted against neural alignment scores relative to the *dense_auto_ - 1_linear_mt* reference model. A linear regression across all 54 models indicates no meaningful association (*R*^2^ = 0.02), suggesting that task-level accuracy in self-motion estimation is not a primary predictor of physiological correspondence in MSTd-like representations.

**Figure 7-1.**
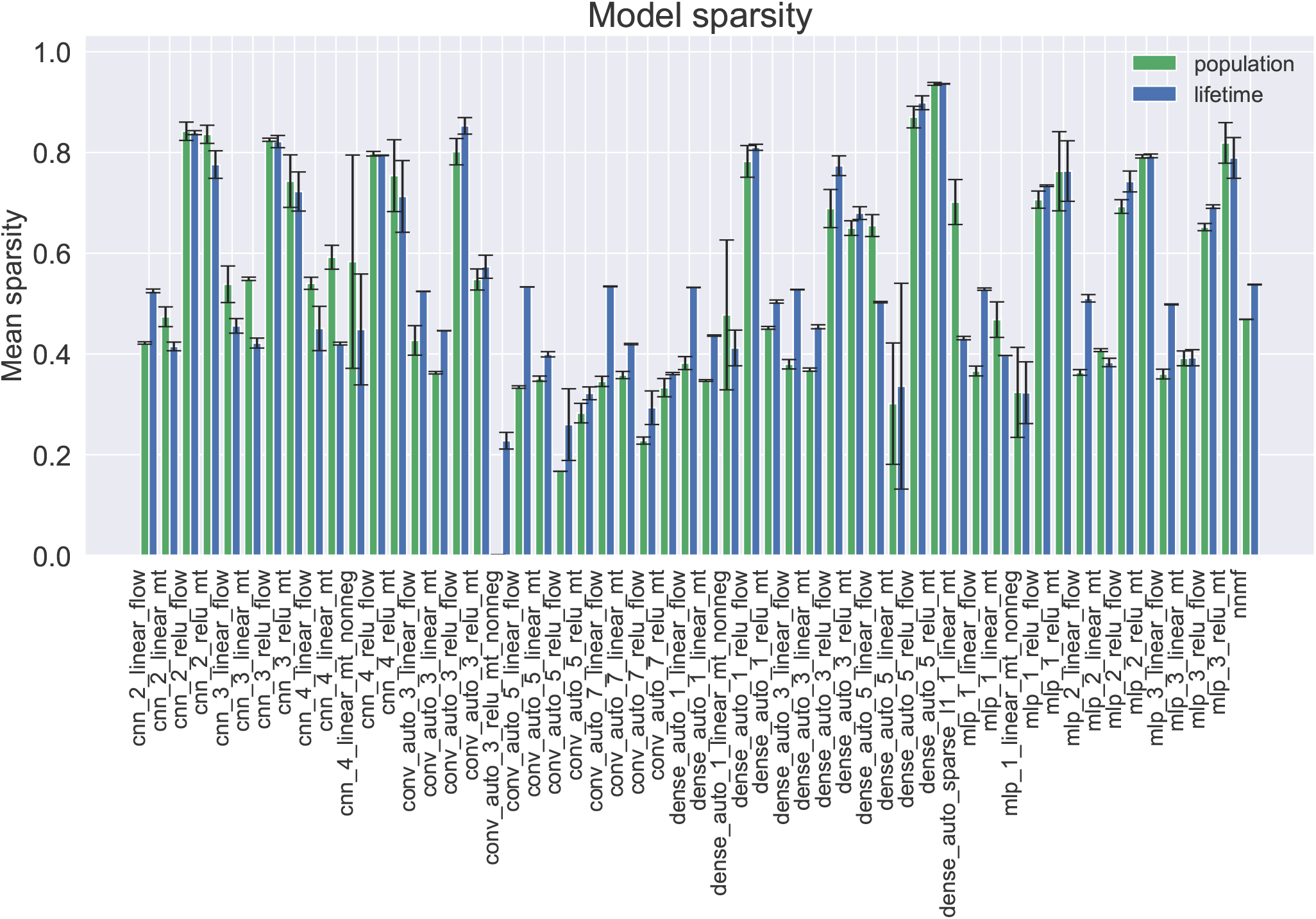
Population and lifetime sparsity across ANN architectures. Sparsity of each model calculated using population (green) and lifetime (blue) metrics. The sparsity metrics range from 0 to 1, where values approaching 1 represent a maximally sparse code and values near 0 reflect a dense representation. Error bars indicate 95% CIs across 25 training repetitions. In most cases, the models that garner the highest sparsity use the ReLU activation function.

